# Scleraxis is required for the growth of adult tendons in response to mechanical loading

**DOI:** 10.1101/2020.03.18.997833

**Authors:** Jonathan P Gumucio, Martin M Schonk, Yalda A Kharaz, Eithne Comerford, Christopher L Mendias

## Abstract

Scleraxis is a basic helix-loop-helix transcription factor that plays a central role in promoting tenocyte proliferation and matrix synthesis during embryonic tendon development. However, the role of scleraxis in the growth and adaptation of adult tendons is not known. We hypothesized that scleraxis is required for tendon growth in response to mechanical loading, and that scleraxis promotes the specification of progenitor cells into tenocytes. We conditionally deleted scleraxis in adult mice using a tamoxifen-inducible Cre-recombinase expressed from the *Rosa26* locus (*Scx*^Δ^), and then induced tendon growth in *Scx*^+^ and *Scx*^Δ^ adult mice via plantaris tendon mechanical overload. Compared to the wild type *Scx*^+^ group, *Scx*^Δ^ mice demonstrated blunted tendon growth. Transcriptional and proteomic analyses revealed significant reductions in cell proliferation, protein synthesis, and extracellular matrix genes and proteins. Our results indicate that scleraxis is required for mechanically-stimulated adult tendon growth by causing the commitment of CD146^+^ pericytes into the tenogenic lineage, and by promoting the initial expansion of newly committed tenocytes and the production of extracellular matrix proteins.

## Introduction

Tendons are made up of an extracellular matrix (ECM) containing primarily type I collagen, as well as other collagens, elastin, and various proteoglycans (1). Tendon ECM is arranged in a hierarchical manner, with densely packed collagen fibers wrapped by layers of basement membrane (1). Tenocytes, or tendon fibroblasts, are the main cell type in tendons and are thought to be responsible for the production, organization, and maintenance of the tendon ECM (2). Tendons are surrounded by an outermost basement membrane called the epitenon, which provides blood and nerve supply to the tendon (1, 3). The organization of the ECM allows the tendon to properly transmit forces from muscle to bone and allow for locomotion, and to respond to mechanical stimuli (4, 5). Mechanical loading can increase tendon cross sectional area (CSA) up to 30% (6) and improve tendon mechanical properties (5, 7), but less is known about the cellular and molecular mechanisms behind tendon growth and adaptation in adult animals.

Scleraxis (*Scx*) is a basic helix-loop-helix (bHLH) transcription factor that is regulated by TGFβ signaling and is required for the proper embryonic development of tendons (8, 9). Scleraxis is the earliest detectible marker for differentiated tendon cells (10), and activates downstream tendon differentiation genes such as type I α-1 chain collagen 1 (*Col1a1*), mohawk (*Mkx*) and tenomodulin (*Tmnd*) through its interaction with the bHLH transcription factor E47, among others (11-13). Scleraxis is expressed through birth and the early postnatal period, but beyond 3 months of age in mice, scleraxis expression in tendons is limited to the epitenon (6). Loss of scleraxis results in the loss or severe disruption of long tendons throughout the body (8). In adult animals, mechanical loading increases scleraxis expression and promotes the proliferation of scleraxis-expressing cells in the epitenon, which correlates to increased type I collagen levels and increased CSA of the tendon (6, 14, 15), while immobilization reduces scleraxis levels (16). Although scleraxis is required for proper embryonic tendon development, it is not known whether scleraxis is required for growth of tendons in adult animals.

Pericytes, or perivascular stem cells, are a population of multipotent stem cells surrounding small blood vessels and are capable of differentiation into many types of mesenchymal cells, including fibroblasts (17). Tendon pericytes express the marker CD146 (*Mcam*) (15, 18, 19). Most of the tenocytes within tendon of adult animals appear to be terminally withdrawn from the cell cycle, with low rates of cell proliferation observed in homeostatic and mechanically loaded tendons (14). Since pericytes are thought to be a progenitor cell population in adult tendons (15, 18, 19), and scleraxis is required for the proper embryonic development of tendons, the objective of this study was to determine the role of scleraxis in adult tendon growth. We hypothesized that scleraxis is required for proper tendon growth in adult animals, and that scleraxis promotes the commitment of pericytes into the tendon lineage. To test this hypothesis, we conditionally deleted scleraxis in adult animals using *Rosa26*^CreERT2/CreERT2^:*Scx*^flox/flox^ mice, and induced tendon growth via mechanical overload of the plantaris tendon. Additionally, we performed a series of *in vitro* experiments with pericytes and tenocytes to gain further mechanistic insight into the observed structural and transcriptional changes in mechanically stimulated tendons that lack scleraxis.

## Results

### Overview

To study the role of scleraxis in adult tendon growth, we generated *Rosa26*^CreERT2/CreERT2^:*Scx*^flox/flox^ mice to allow for the inactivation of scleraxis upon treatment with tamoxifen (referred to as *Scx*^Δ^ mice), while *Rosa26*^CreERT2/CreERT2^:*Scx*^+/+^ mice maintain the expression of *Scx* after tamoxifen treatment (referred to as *Scx*^+^ mice). An overview of the *Rosa26*^CreERT2^, *Scx*^+^, *Scx*^flox^, and *Scx*^Δ^ alleles is provided in Figure 1A. Following tamoxifen treatment, we induced a supraphysiological overload of the plantaris tendons of *Scx*^+^ and *Scx*^Δ^ mice by synergist ablation, and then analyzed tendons at either 7D or 14D after surgery (Figure 1B-C). We also isolated cells from *Scx*^+^ and *Scx*^Δ^ mice to further mechanistically study the effects of scleraxis on the behavior of tenocytes *in vitro*. To verify efficient deletion of scleraxis, we isolated DNA and RNA from tendons of animals treated with tamoxifen, and cultured cells treated with 4-hydroxytamoxifen (4HT). There was a 97% and 93% reduction in *Scx* exon 1 abundance in 7D and 14D plantaris tendons, respectively (Figure 1D). Cultured tenocytes of *Scx*^Δ^ mice had a 99% reduction in exon 1 abundance compared to *Scx*^+^ tenocytes (Figure 1F). RNAseq data also revealed a significant downregulation in *Scx* transcript levels in plantaris tendons and cultured cells (Figure 1E, G). These data confirm the efficient conditional deletion of scleraxis in the models used in this study.

**Figure 1.**
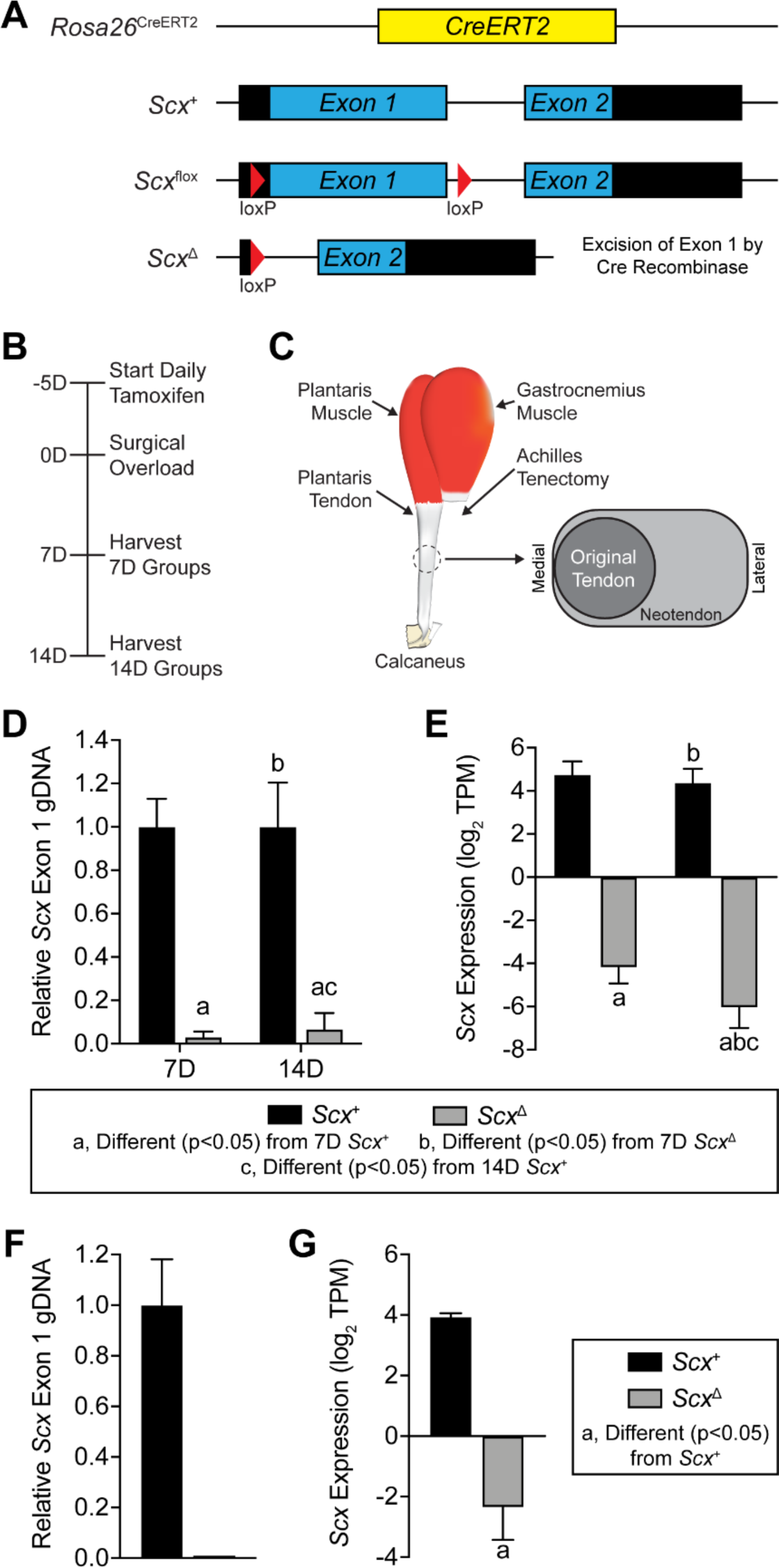
Experimental overview. (A) Overview of the alleles used to perform targeted deletion of scleraxis in this study, including the constitutive CreERT2 (*Rosa26*^CreERT2^), scleraxis wild type (*Scx*^+^), scleraxis floxed (*Scx*^flox^), and scleraxis loss-of- function (*Scx*^Δ^) alleles. (B) Timeline of tamoxifen treatment, synergist ablation/plantaris growth procedure, and tissue harvest. Tamoxifen was delivered on a daily basis beginning five days prior to surgery, and continued through tissue harvest. (C) Overview of synergist ablation/plantaris growth procedure. A large segment of the Achilles tendon is carefully removed from the animal, resulting in compensatory hypertrophy of the synergist plantaris muscle and tendon. A neotendon area of new tendon extracellular matrix forms around the original tendon, growing in a lateral direction towards the area where the Achilles was previously located. (D, F) Relative genomic DNA (gDNA) abundance of exon 1 of scleraxis to exon 2, as measured using qPCR, in (D) plantaris tendons of *Scx*^+^ and *Scx*^Δ^ mice either 7D or 14D after synergist ablation, or (F) in cultured *Scx*^+^ or *Scx*^Δ^ tenocytes. (E, G) Expression of *Scx*, measured using RNAseq and quantified as transcripts per kilobase million (TPM) reads, in (E) plantaris tendons of *Scx*^+^ and *Scx*^Δ^ mice either 7D or 14D after synergist ablation, or (F) in cultured *Scx*^+^ or *Scx*^Δ^ tenocytes. Values are (D, F) mean±CV or (E, G) mean±SD. (D-E) Differences between groups were tested using a two-way ANOVA: a, significantly different (p<0.05) from 7D *Scx*^+^; b, significantly different (p<0.05) from 7D *Scx*^Δ^; c, significantly different (p<0.05) from 14D *Scx*^+^. (F-G) Differences between groups were tested using a t-test: a, significantly different (p<0.05) from *Scx*^+^ tenocytes. N=4 per group.

### Scleraxis deletion impairs tendon growth

Following synergist ablation, *Scx*^+^ mice demonstrated an increase in plantaris tendon cross section area (CSA) through the formation of neotendon matrix, while this response was blunted in *Scx*^Δ^ mice (Figure 2A). The original tendon CSA and cell density did not change between groups (Figure 2B, C). However, at 7 days *Scx*^Δ^ mice had a 60% reduction in neotendon CSA which resulted in a 36% smaller total tendon area (Figure 2B). At 14 days, *Scx*^Δ^ mice had a 65% decrease in neotendon CSA and a 47% decrease in overall tendon CSA compared to *Scx*^+^ mice (Figure 2B). For cell density, at 7 days there was a 34% decrease in the neotendon and total tendon of *Scx*^Δ^ mice (Figure 2C). Collagen fibril size distributions were significantly different between the original tendon and neotendon of *Scx*^+^ and *Scx*^Δ^ mice at 7 and 14 days, with *Scx*^Δ^ mice generally demonstrating a greater proportion of larger collagen fibrils (Supplemental Figure 1).

**Figure 2.**
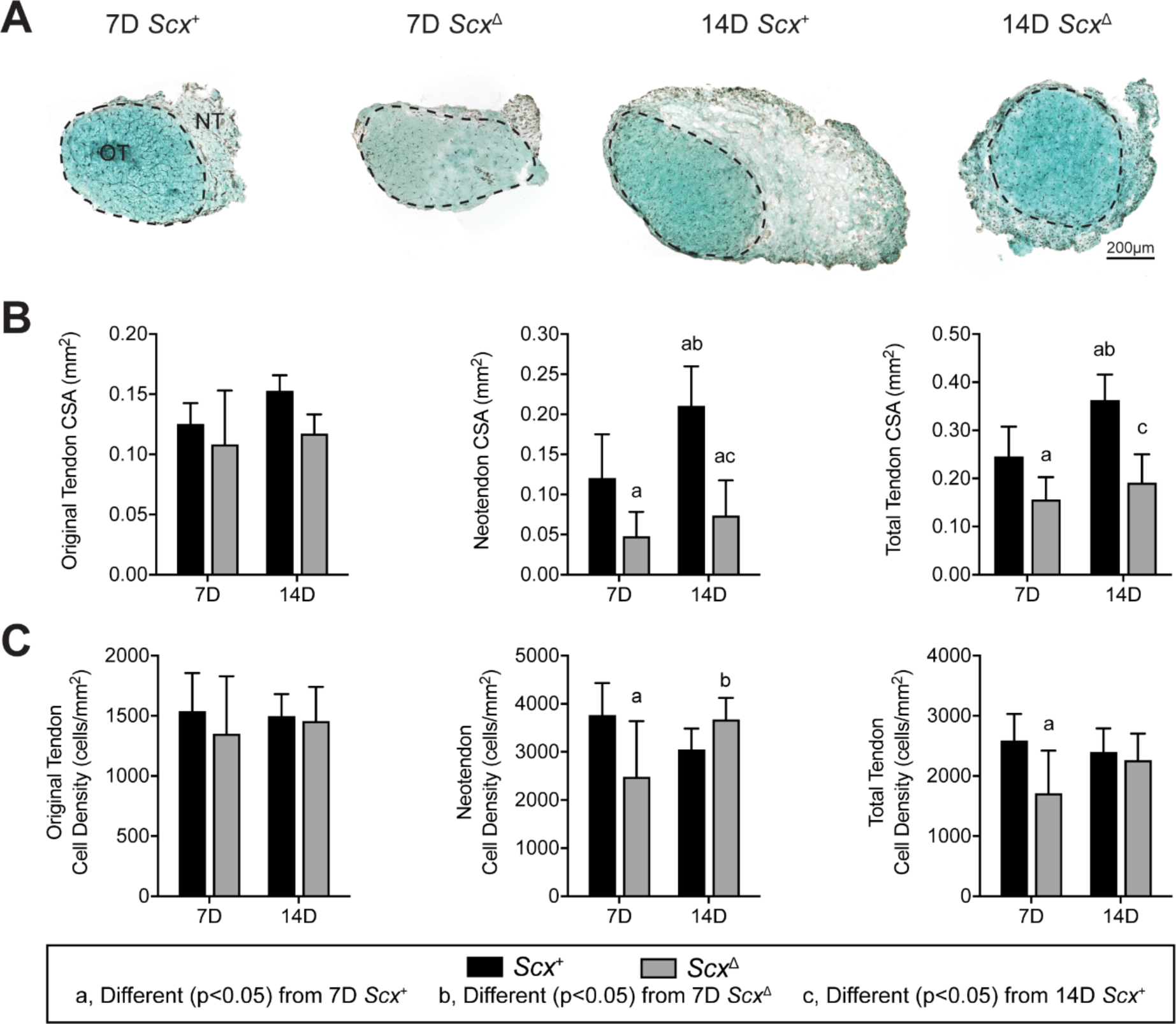
Effect of scleraxis deletion on tendon growth. (A) Representative fast green stained cross section histology from *Scx*^+^ and *Scx*^Δ^ mice at either 7D or 14D after synergist ablation/plantaris growth procedure demonstrating general morphology and cell density. The original tendon (OT) and neotendon (NT) are indicated by the hashed line. (A) Scale bar for all images is 200µm. Quantification of original tendon, neotendon, and total tendon (B) cross-sectional area (CSA) and (C) cell density. Values are mean±SD. Differences between groups were tested using a two-way ANOVA: a, significantly different (p<0.05) from 7D *Scx*^+^; b, significantly different (p<0.05) from 7D *Scx*^Δ^; c, significantly different (p<0.05) from 14D *Scx*^+^. N≥4 mice per group.

### Scleraxis deletion increases pericyte density

Using immunohistochemistry, we measured the abundance of CD146^+^ pericytes within the neotendon of mechanically overloaded plantaris tendons (Figure 3A). Compared to *Scx*^Δ^ mice, there was an approximate two-fold increase in the percentage of pericytes in the neotendons of *Scx*^+^ mice at each time point (Figure 3B). As CD146^+^ pericytes are thought to be a progenitor cell for tenocytes, and given the size differences between *Scx*^+^ and *Scx*^Δ^ mice, we determined whether the percentage of pericytes in the neotendon correlated with tendon size. When all time points and genotypes were combined, there was a negative correlation (r=-0.58) between neotendon CSA and pericyte cell abundance (Figure 3C).

**Figure 3.**
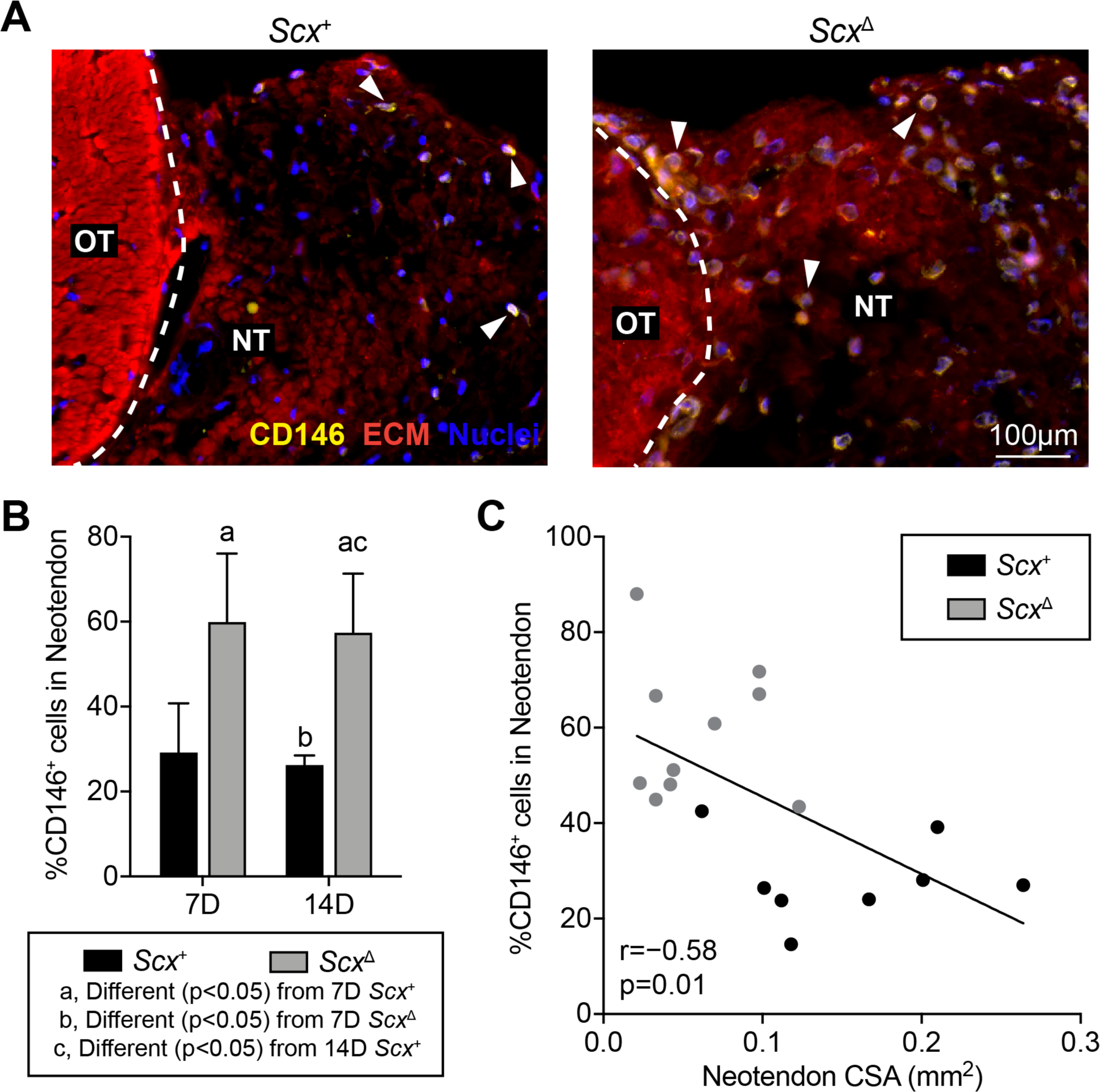
Effect of scleraxis deletion on pericyte density. (A) Representative immunofluorescent histology demonstrating the presence of CD146+ pericytes in tendons of *Scx*^+^ and *Scx*^Δ^ mice 14D after synergist ablation/plantaris growth procedure. The original tendon (OT) and neotendon (NT) are indicated by the hashed line. CD146, yellow; extracellular matrix (ECM), red; nuclei, blue. Scale bar for both images is 100µm. (B) Quantification of the CD146+ pericytes as a percentage of total cells in the neotendon in *Scx*^+^ and *Scx*^Δ^ mice at 7D or 14D. Values are mean±SD. Differences between groups were tested using a two-way ANOVA: a, significantly different (p<0.05) from 7D *Scx*^+^; b, significantly different (p<0.05) from 7D *Scx*^Δ^; c, significantly different (p<0.05) from 14D *Scx*^+^. (C) Percentage of CD146+ pericytes across all 7D and 14D *Scx*^+^ and *Scx*^Δ^ mice as a function of the neotendon CSA, with the Pearson r and p-value presented. N≥4 mice per group.

### Scleraxis deletion impacts the proteome of tendons 14 days after synergist ablation

As we observed noticeable differences between the size of tendons from mechanically loaded *Scx*^+^ and *Scx*^Δ^ mice, we performed mass spectrometry-based label-free proteomics to look at changes in tendon matrix composition. Principal component analyses (PCA) demonstrated differences between overloaded groups and the control non-overloaded group, as well as some non-overlapping areas between the *Scx*^+^ and *Scx*^Δ^ groups (Figure 4A). In total, 313 proteins out of the 453 that were detected were found to be significantly different between *Scx*^+^ and *Scx*^Δ^ mice for at least one time point, with 298 of these proteins being significantly different at 14 days. We report select proteins in Figure 4B, with the entire dataset available in Supplemental Table 1. At 7 days, the matrix proteins chondroadherin (Chad) and cartilage intermediate layer protein (Cilp) were reduced in *Scx*^Δ^ tendons, which was also noted at 14 days. Additionally, at 14 days several collagens, including Col3a1, Col5a1, Col11a1, Col12a1, were reduced in *Scx*^Δ^ tendons compared to *Scx*^+^ tendons. Proteins involved in extracellular matrix production, remodeling, and mechanotransduction, such as asporin (Aspn), fibronectin (Fn1), lysyl oxidase (Lox), matrix metalloproteinase 2 (Mmp2), osteoglycin (Ogn), thrombospondins 1 and 4 (Thbs1, Thbs4), and tenascin C (Tnc) were lower in tendons of *Scx*^Δ^ mice (Figure 4B). Ribosomal proteins Rplp0, Rpl10, and Rpl35, eukaryotic translation elongation factors Eef1g and Eef2, and mechanotransduction proteins cofilin 1 (Cfl1), gelsolin (Gsn), Ras homolog family member C (Rhoc), and vimentin (Vim) were also less abundant in *Scx*^Δ^ tendons at 14 days (Figure 4B).

**Table 1.** Pathway enrichment analysis. P-values of select affected or differentially regulated pathways or biological functions that were identified using Ingenuity Pathway Analysis. ND, not significantly different.

| Pathway | 7D Scx<sup>+</sup> to<br>7D Scx<sup>Δ</sup> | 14D Scx<sup>+</sup> to<br>14D Scx<sup>Δ</sup> | Cultured<br>Tenocytes Scx<sup>Δ</sup><br>to Scx<sup>+</sup> |
| --- | --- | --- | --- |
| Actin Cytoskeleton Signaling | ND | ND | 2.45E-02 |
| AMPK Signaling | 8.87E-03 | ND | 3.15E-03 |
| BMP Signaling | ND | ND | 3.55E-02 |
| ERK/MAPK Signaling | ND | 3.14E-02 | 1.44E-05 |
| IGF1 Signaling | 1.26E-02 | 7.24E-04 | 1.02E-06 |
| IL1 Signaling | 4.37E-03 | 1.26E-02 | 1.35E-05 |
| IL6 Signaling | 9.77E-10 | 1.58E-03 | 4.11E-11 |
| Inhibition of MMPs | 1.23E-02 | 3.02E-02 | 7.76E-04 |
| NFκB Signaling | 2.34E-02 | ND | ND |
| p38 MAPK Signaling | 3.76E-02 | 1.59E-03 | 5.53E-10 |
| PI3K Signaling | 5.64E-03 | 1.61E-04 | 1.18E-04 |
| PDGFB Signaling | ND | 1.10E-03 | 5.11E-04 |
| RhoA Signaling | ND | 3.61E-03 | 8.91E-03 |
| STAT3 Pathway | 9.43E-26 | 1.19E-03 | 5.61E-08 |
| TGFβ Signaling | 4.61E-03 | 8.49E-06 | 4.34E-08 |
| VEGFA Signaling | ND | 2.26E-02 | 1.27E-05 |
| Wnt Signaling | ND | 4.00E-02 | 4.03E-02 |

**Figure 4.**
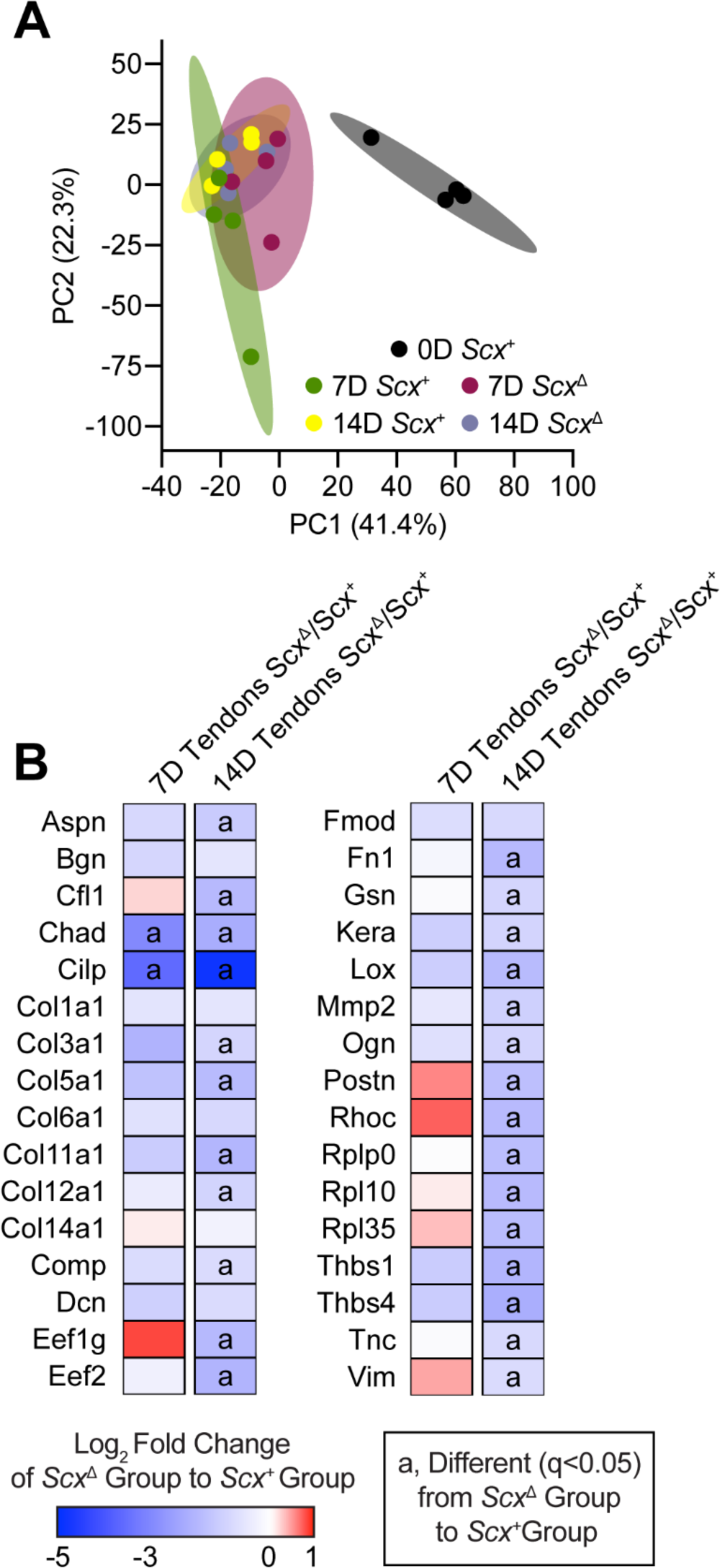
Effect of scleraxis deletion on the proteome of tendons. (A) Principal component (PC) analysis of mass spectroscopy-based proteomics data of tendons from control non-overloaded (0D) *Scx*^+^ tendons, as well as from *Scx*^+^ and *Scx*^Δ^ mice at 7D or 14D after synergist ablation/plantaris growth procedure. (B) Heatmaps demonstrating the log2 fold change in selected proteins between *Scx*^+^ and *Scx*^Δ^ tendons at 7D and 14D. Differences between groups were tested using FDR-corrected t-tests: a, different (q<0.05) between *Scx*^+^ and *Scx*^Δ^ groups at a given time point. N=4 mice per group.

### Scleraxis deletion impacts the transcriptome of tendons and cultured tenocytes

We then performed RNA sequencing in order to analyse transcriptional changes in overloaded plantaris tendons from *Scx*^+^ and *Scx*^Δ^ mice, and in cultured tenocytes. Principal component analyses demonstrated both unique and overlapping regions for tendons of *Scx*^+^ and *Scx*^Δ^ mice at 7 and 14 days, and all four groups were different from control, non-overloaded tendons (Figure 5A). More divergence in genotypes was observed in cultured tenocytes (Figure 5B). Seven and fourteen days after synergist ablation, there were 679 and 177 transcripts, respectively, that were significantly different and at least 50% differentially regulated (Figure 5D-E). *In vitro*, there were 680 genes that were significantly different between *Scx*^+^ and *Scx*^Δ^ tenocytes (Figure 5F).

**Figure 5.**
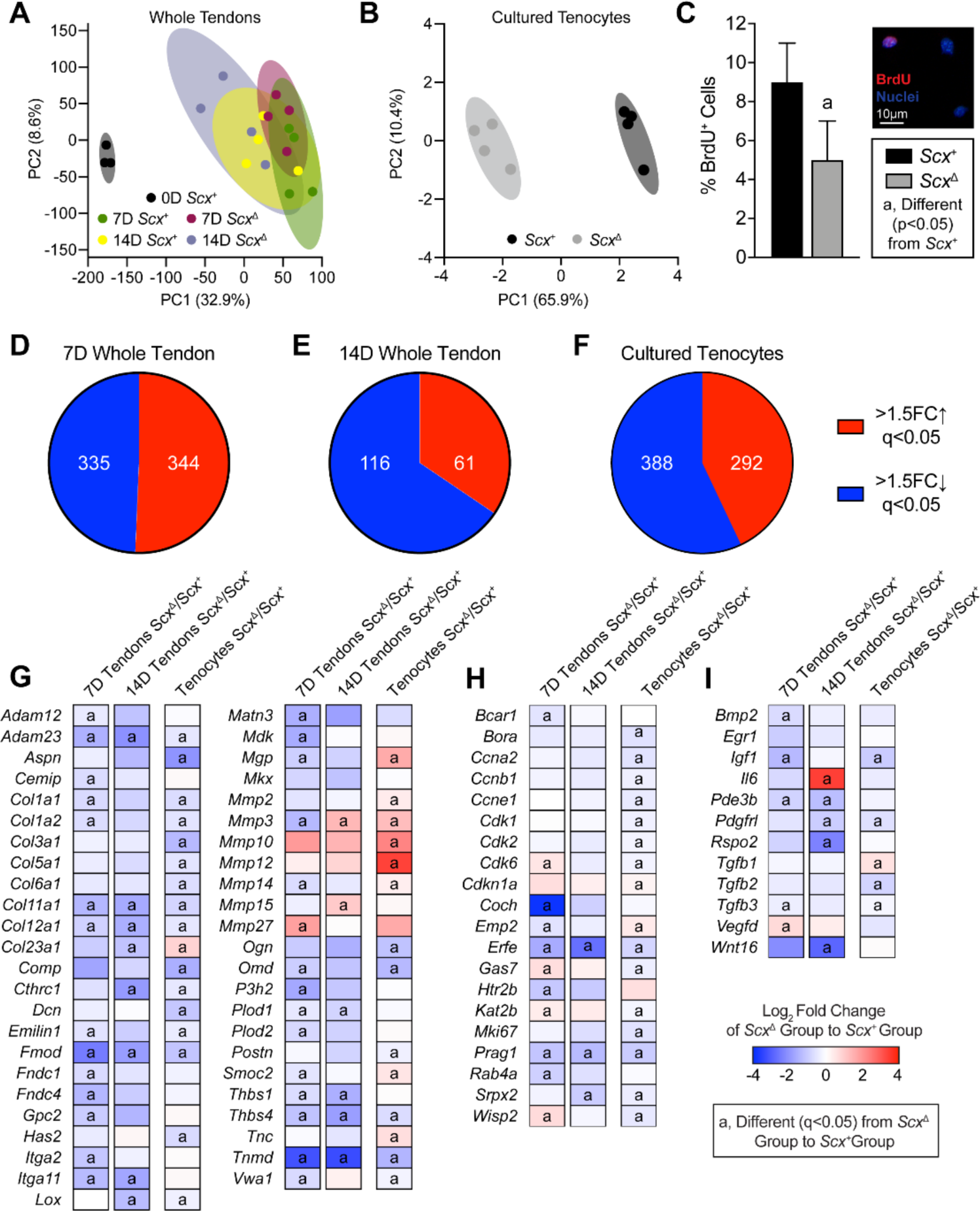
Effect of scleraxis deletion on transcriptional changes of whole tendons and cultured tenocytes. Principal component (PC) analysis of RNA-sequencing data of (A) tendons from control non-overloaded (0D) *Scx*^+^ tendons, as well as from *Scx*^+^ and *Scx*^Δ^ mice at 7D or 14D after synergist ablation/plantaris growth procedure, and (B) from cultured *Scx*^+^ and *Scx*^Δ^ tenocytes. (C) Proliferating tenocytes as quantified by BrdU+ nuclei (red) as a percentage of total nuclei (blue) in *Scx*^+^ and *Scx*^Δ^ tenocytes. Representative BrdU- and BrdU+ cells are shown in the inset. Scale bar is 10µm. (D-F) The number of genes that with a > 1.5-fold upregulation (red) or a > 1.5-fold downregulation (blue), and with q-values < 0.05, in *Scx*^Δ^ tendons compared to *Scx*^+^ tendons at (D) 7D or (E) 14D after synergist ablation/plantaris growth procedure, or in (F) cultured *Scx*^Δ^ tenocytes compared to *Scx*^+^ tenocytes. (G-I) Heatmaps demonstrating the log2 fold change in selected genes from RNA-sequencing that are (G) components or regulators of the extracellular matrix or tendon differentiation, (H) involved in cell migration or proliferation, or (I) growth factors, cytokines, and signaling molecules. The fold change value is displayed for *Scx*^Δ^ relative to *Scx*^+^ group. (C) Values are mean±SD. Differences between *Scx*^+^ and *Scx*^Δ^ groups were tested using a t-test. (G-I) Differences between groups were tested using DESeq2: a, different (q<0.05) between *Scx*^+^ and *Scx*^Δ^ groups at a given time point for whole tendons, or for cultured tenocytes. (A,D,E,G-I) N≥3 tendons and (B,C,F-I) N=6 replicates per tenocyte group.

Pathway enrichment analysis was performed for both experiments to determine signaling pathways predicted to be different between groups (Table 1). Several of the pathways identified were involved with growth and differentiation, cytoskeletal signaling, and ECM production. Based on these findings, and from genes that are known to be important for tenocyte proliferation and tenogenesis, we selected several genes for reporting in the manuscript, with the full dataset available on NIH GEO. We also performed qPCR validation for a select set of genes of interest identified from RNAseq data, and generally observed similar patterns of fold change differences between tendons of *Scx*^+^ and *Scx*^Δ^ mice (Table 2). Many genes in the whole tendon RNAseq data were also differently expressed in the cultured cells.

**Table 2.**
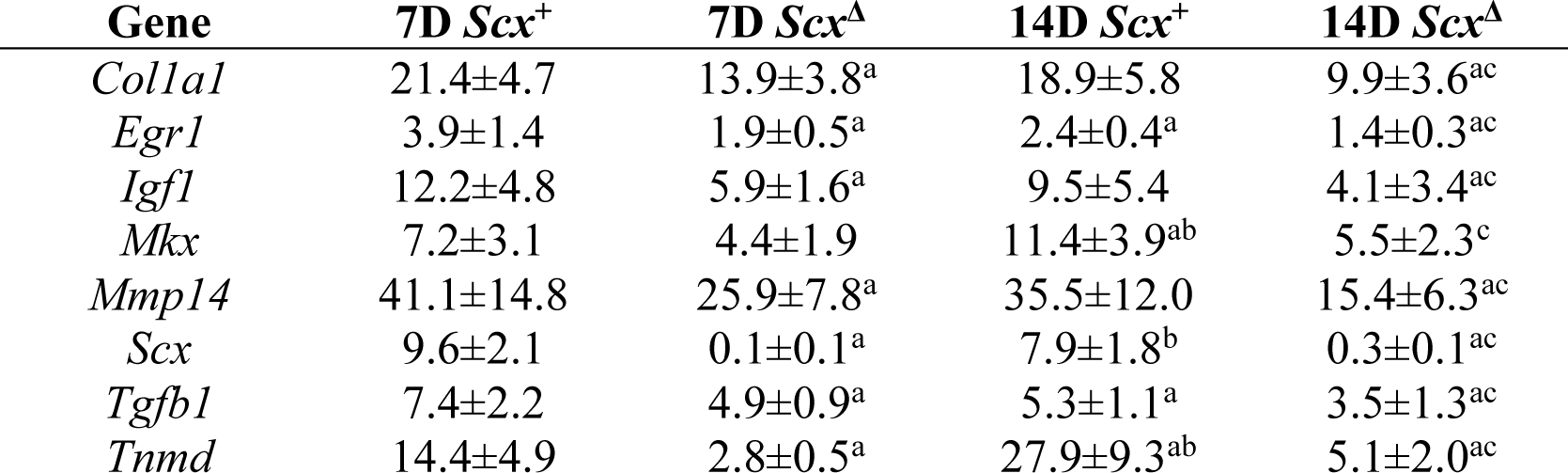
Gene expression data. Expression of genes as measured by qPCR. Expression is normalized to 0D non-overloaded control *Scx*^+^ mice. Values are mean±SD. Differences were tested with a two-way ANOVA (α=0.05): a, different (p<0.05) from 7D *Scx*^+^; b, (different (p<0.05) from 7D *Scx*^Δ^; c, different (p<0.05) from 14D *Scx*^+^. ND, not detected.

Expression of components or regulators of the ECM, including the collagens *Col1a1, Col1a2, Col11a1*, and *Col12a*, small leucine-rich proteoglycans *Aspn, Dcn, Fmod*, and *Omd*, and proteases *Adam12* and *Adam23* were downregulated in *Scx*^Δ^ whole tendons and cultured tenocytes, while some matrix degradation genes, such as *Mmp10* and *Mmp12*, were upregulated (Figure 5G). There were also genes like *Col23a1, Mgp, Mmp14* and *Smoc2* that displayed differential expression patterns between whole tendons and cultured tenocytes (Figure 5G).

Genes involved in cell proliferation and migration were different between genotypes mostly at the 7D time point (Figure 5H). Many cyclins and cyclin dependent kinases, such as *Ccna2*, and *Cdk1, 2* and *6*, as well as the cell proliferation marker *Mki67*, were significantly downregulated in the cultured *Scx*^Δ^ tenocytes compared to *Scx*^+^ tenocytes, with similar patterns in whole tendon data (Figure 5H). Consistent with this, cell proliferation as measured with BrdU incorporation, was reduced by 44% in *Scx*^Δ^ tenocytes compared to *Scx*^+^ tenocytes (Figure 5C). Several growth factors and signaling molecules were also reduced in *Scx*^Δ^ tendons and cells, including *Igf1, Pdgfrl, Tgfb2*, and *Tgfb3* (Figure 5I).

### Scleraxis deletion reduces the ability of pericytes to commit to the tenogenic lineage

As we observed an accumulation of CD146^+^ pericytes in the neotendon of *Scx*^Δ^ mice compared to *Scx*^+^ mice, we sought to evaluate the effect of scleraxis deletion on the commitment of pericytes to the tenogenic lineage *in vitro*. We first wanted to identify culture conditions that would result in pericyte differentiation into tenocytes. We found that culturing pericytes on type I collagen coated plates, which is similar to the mature tendon matrix, induced morphological changes in cells to make them appear like tenocytes and express *Col1a1, Mkx, Scx*, and *Tnmd* (Supplemental Figure 2A-B). However, culturing pericytes on tissue culture plastic alone, or on basement membrane-derived Matrigel, which is similar to the perivascular matrix of tendon, did not induce markers of tenogenesis (Supplemental Figure 2A-B).

After establishing that pericytes isolated from tendons and placed on type I collagen appear to enter the tenogenic lineage, we sought to determine how the deletion of scleraxis would impact this process. *Scx*^+^ and *Scx*^Δ^ CD146^+^ pericytes were cultured on type I collagen substrates for 1, 3 and 5 days, and qPCR was performed (Figure 6). As expected, *Scx* was undetectable in the *Scx*^Δ^ cells at all time points, but increased at 3 and 5 days in the *Scx*^+^ cells (Figure 6). Relative expression of *CD146* decreased over time in the *Scx*^+^ group while it increased in the *Scx*^Δ^ group (Figure 6). *Col1a1, Mkx* and *Tnmd* expression significantly increased over time in the *Scx*^+^ cells, and to a much greater extent than in *Scx*^Δ^ cells (Figure 6).

**Figure 6.**
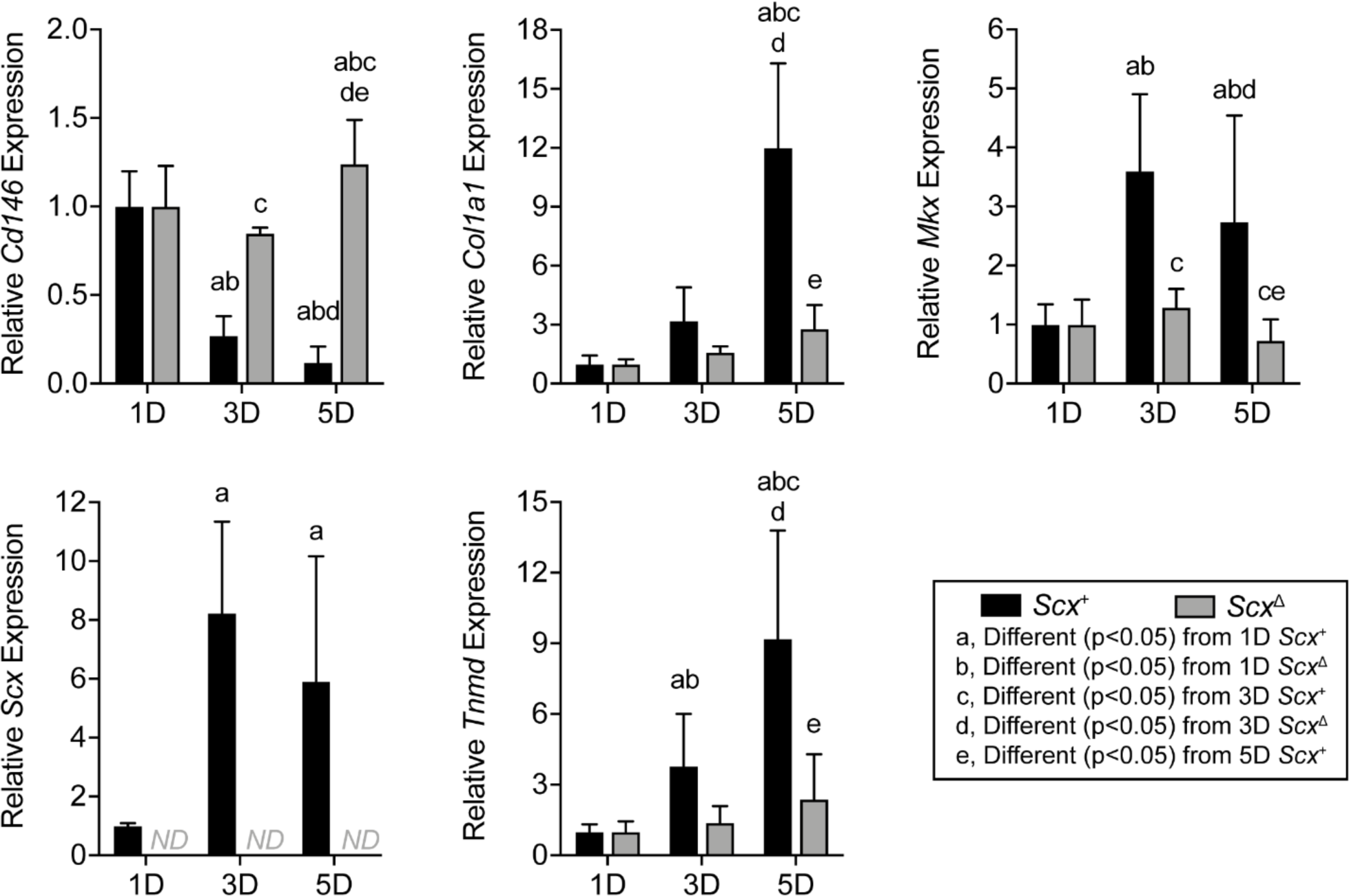
Effect of scleraxis deletion on gene expression in cultured pericytes. Gene expression (*Cd146, Col1a1, Mkx, Scx*, and *Tnmd*) of CD146+ MACS-sorted pericytes from *Scx*^+^ and *Scx*^Δ^ mice cultured on type I collagen gels for 1, 3 or 5 days. Values are mean±CV. Differences between groups were tested using a two-way ANOVA: a, significantly different (p<0.05) from 1D *Scx*^+^; b, significantly different (p<0.05) from 1D *Scx*^Δ^; c, significantly different (p<0.05) from 3D *Scx*^+^; d, significantly different (p<0.05) from 3D *Scx*^Δ^; e, significantly different (p<0.05) from 5D *Scx*^+^. Values are mean±SD. N=6 replicates per group.

## Discussion

While the cellular components and genetic program that are responsible for the initial embryonic formation and elongation of tendons are well understood (20-23), less is known about growth of tendon tissue in adult animals. Using the synergist ablation model, we previously identified the formation of a neotendon matrix that develops around the original tendon matrix, and that this new matrix was populated with proliferative, scleraxis-expressing cells (14). The aim of our study was to identify whether scleraxis was required for tendon adaptation to growth stimuli. To study tendon growth, we used the synergist ablation mouse model, in which the Achilles tendon is surgically removed resulting in a supraphysiological growth stimulus to the plantaris tendon and muscle (15, 24-26). A neotendon matrix consisting of immature collagen and other ECM proteins forms around the plantaris tendon, and over a month period of time this ECM matures and resembles the original tendon matrix (15). Our results indicate that scleraxis is required for the proper expansion, formation, and remodeling of tendon ECM during mechanical loading, and that growth defects in scleraxis-deficient tendons appear to occur in part due to a reduction in the ability of pericytes to differentiate into the tendon cell lineage.

The existing tenocytes within adult tendons appear to be mostly postmitotic cells that do proliferate in response to mechanical loading (14, 27, 28). Increased numbers of tenocytes that occur in response to growth stimuli have been postulated to occur through differentiation of tissue resident progenitor cells into tenocytes (15, 18, 19, 28, 29). TGFβ signaling plays an important role in the proper embryonic development of tendons by regulating scleraxis and other signaling events (9, 20). Pathway enrichment analysis in the current study demonstrated a potentially important role for TGFβ signaling and other pathways such as IL1, IL6, p38 MAPK, and STAT3 in postnatal tendon growth. Several studies have suggested that one of the populations of progenitor cells in adult tendon tissue are CD146^+^ pericytes, located adjacent to tendon vasculature (15, 18, 19, 30). In a rat model of patellar tendon injury and repair, these cells participate in tenogenic differentiation when stimulated with CTGF (18). In the patellar tendon, approximately 0.8% of cells are CD146^+^ (18), which is consistent with the relatively low amount of vasculature within homeostatic adult tendon (3). Previous studies in our laboratory using the synergist ablation model in rats showed that these cells appear to migrate from their niche in the vasculature and proliferate within the neotendon (15). In the current study, despite smaller tendon CSAs in *Scx*^Δ^ mice, the relative abundance of CD146^+^ pericytes was twice as high in the neotendon of *Scx*^Δ^ mice compared to *Scx*^+^ mice, and the percentage of pericytes was negatively correlated with tendon CSA. When pericytes were cultured in tenogenic conditions, deleting scleraxis prevented the ability of these cells to express tenogenic markers. When taken together, these *in vivo* and *in vitro* findings indicate that scleraxis is required for the commitment of pericytes into the tenogenic lineage. Further, the higher percentage of CD146^+^ pericytes in the neotendon of *Scx*^Δ^ mice likely occurs due to the failure of these cells to properly differentiate into tenocytes. Alternatively, scleraxis could inhibit pericyte proliferation *in vivo*. While we did not directly measure pericyte proliferation in tendons, the *in vitro* data suggests that a reduction in the specification or differentiation of pericytes into the tenogenic lineage is more likely to be responsible for the greater abundance of pericytes observed in *Scx*^Δ^ mice.

During early postnatal development, committed tenocytes (sometimes referred to as tenoblasts) are able to proliferate for a period of time (28), and a similar proliferative capacity likely exists for newly committed tenocytes that arise from CD146^+^ progenitors. This is further supported by previous work that demonstrated scleraxis-expressing cells in the neotendon of mechanically stimulated tendons also display uptake of the cell proliferation marker EdU (14). In the current study, several genes that are involved in cell proliferation were downregulated in *Scx*^Δ^ tendons or cells, including numerous cyclins and cyclin dependent kinases, as well as the cell proliferation marker *Mki67*. Further, *Scx*^Δ^ tenocytes displayed a proliferation rate that was nearly half that of *Scx*^+^ tenocytes. These results indicate that scleraxis also induces postnatal tenocyte proliferation.

Scleraxis appears to be important in embryonic tendon development, in part, by directing the expression of type I collagens, tenomodulin, and fibronectin (12, 13, 31-33). Mechanical loading increases the expression of scleraxis, type I and III collagen, and tenomodulin in tendon (6, 14, 34, 35), and our transcriptomic analyses demonstrated that expression of components or regulators of the ECM, including *Col1a1, Mkx* and *Tnmd*, were found to be reduced in the absence of scleraxis in whole tendons and in cultured tenocytes. Proteoglycans and glycoproteins that compose the tendon ECM, such as asporin, fibronectin, osteoglycin, thrombospondins 1 and 4, and tenascin C were lower in tendons of *Scx*^Δ^ mice at 14 days, although no change in the expression of many genes involved with heparin sulfate, chondroitin sulfate, or hyaluronic acid synthesis were observed in whole tendons. Differences in MMPs, FACIT collagens, and other factors that regulate type I collagen fibril diameter may also explain the apparent differences in fibril size distributions in electron micrographs. IGF1 signaling can induce protein synthesis in tenocytes (26), and *Igf1* expression was downregulated in scleraxis-deficient mice and cultured cells, along with reduced ribosomal proteins and translation elongation factor proteins in tendons of *Scx*^Δ^ mice. The growth deficiency of *Scx*^Δ^ tendons was most pronounced at 14 days, and proteomics analyses at this time point identified a reduction in several collagens and proteoglycans. These results indicate that, similar to embryonic development, scleraxis plays an important role in adult tendon growth by directing the expression of ECM proteins. When combined with other findings in this manuscript, we propose a model of mechanically-stimulated adult tendon growth whereby scleraxis plays a central role in the commitment of CD146^+^ pericytes to the tenogenic lineage, the initial expansion of newly committed tenocytes, and the production of ECM proteins (Figure 7).

**Figure 7.**
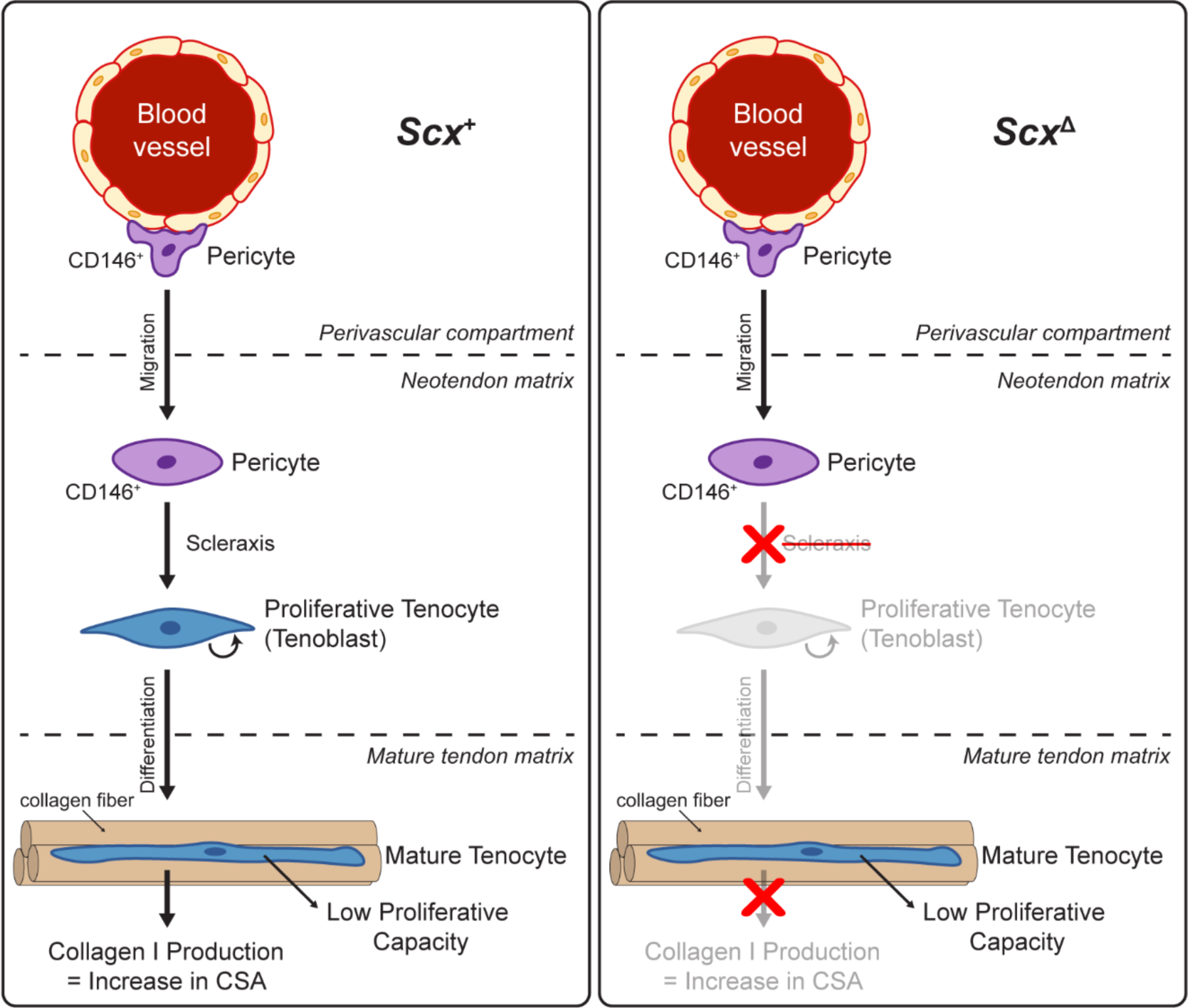
Proposed model of scleraxis role in adult tendon growth response to mechanical loading. In wild-type (left) scleraxis controls the differentiation of CD146+ pericytes into a tenogenic lineage. When scleraxis is deleted (right), CD146+ cells still migrate to the neotendon but are unable to differentiate and do not participate in ECM synthesis.

Despite providing substantial new insight into the role of scleraxis in adult tendon adaptation, this study is not without limitations. We only studied male mice, as we previously demonstrated a general lack of differences between the Achilles tendons of male and female mice (36), and we anticipate the results are likely applicable to both sexes. Results from the plantaris tendon may not be applicable to all tendons, as different tendons display extensive divergence in their function and transcriptional profiles (37). Only two time points were used following overload to study scleraxis function. These time points were selected due to high levels of scleraxis expression in a previous study using the same tendon growth model (14), but additional insight would likely be gained from studying long term changes in scleraxis-deficient tendon growth. While we achieved efficient knock down of scleraxis in whole tendons and cultured tenocytes, there are likely some cells in which scleraxis was not deleted. Although we and others demonstrated that CD146^+^ pericytes likely serve as progenitor cells for tenocytes, we did not test this with genetic lineage tracing. We also did not directly determine which genes that scleraxis directly regulates by recruitment or repression of transcription machinery to promoter or enhancer sequences. Immune cells likely contribute to tendon growth, and we did not directly quantify the abundance of these cell populations with histology or flow cytometry. However, in the extended RNAseq dataset we did not observe differences in the expression of the pan-hematopoietic cell marker *Cd45*, macrophage markers (such as *Cd11b, Cd68, Cd163, Cd206*), or T cell markers (*Cd3, Cd4, Cd8*) between tendons of *Scx*^+^ and *Scx*^Δ^ mice. Lastly, while the synergist ablation model is useful in the study of mechanical load-induced tendon growth, the model induces supraphysiological growth that exceeds the loading that is used clinically to induce tendon hypertrophy, such as progressive resistance exercise. There may be a larger inflammatory stimulus in the synergist ablation model compared to resistance exercise, but we think the fundamental mechanisms of tendon growth and ECM synthesis are similar between the two conditions.

Tendons of adult animals display an ability to grow and adapt in response to mechanical loading (5-7), and deficiencies in this process may lead to the development of painful and debilitating tendinopathies (38, 39). We found that scleraxis plays an important role in the growth response of adult tendons to mechanical loading by promoting the commitment and expansion of new tenocytes, as well as the production of tendon ECM components. Scleraxis may therefore be a useful biomarker in assessing the response of rehabilitation interventions to treat tendinopathies. This study also provides further support for the potential therapeutic use of pericytes in the treatment of tendon disorders. While scleraxis appears to be a critical transcription factor for tendon growth throughout the lifespan, additional studies which integrate mechanotransduction with molecular genetics and cell biology will provide further insight into the basic mechanisms of tendon growth and contribute to the development of novel biomarkers of early tendinopathic changes and improved therapies for tendinopathies.

## Methods

### Mice

Wild type C57BL/6J (WT, strain 000664) mice, and *Rosa26*^CreERT2^ mice (strain 008463) in which the tamoxifen-sensitive CreERT2 recombinase enzyme is expressed from the ubiquitous *Rosa26* locus (40) were obtained from the Jackson Laboratory (Bar Harbor, ME, USA). *Scx*^flox/flox^ mice in which loxP sites flank the coding region of exon 1 of scleraxis (8), and transgenic *Scx*GFP mice in which GFP is driven by 4kb of the scleraxis promoter (41), were also used. Genotype of mice was determined by PCR analysis of DNA obtained from tail tendon biopsies. After performing initial crosses between *Rosa26*^CreERT2/CreERT2^ and *Scx*^flox/flox^ mice, we generated *Rosa26*^CreERT2/CreERT2^:*Scx*^flox/flox^ mice to allow for the inactivation of scleraxis upon treatment with 4-hydroxytamoxifen (4HT) *in vitro* or tamoxifen *in vivo* (referred to as *Scx*^Δ^ mice), while *Rosa26*^CreERT2/CreERT2^:*Scx*^+/+^ mice maintain the expression of *Scx* after 4HT or tamoxifen treatment (referred to as *Scx*^+^ mice). An overview of the *Rosa26*^CreERT2^, *Scx*^+^, *Scx*^flox^, and *Scx*^Δ^ alleles is provided in Figure 1A. Male mice 4-6 months were used in this study.

### Scleraxis inactivation in mice

*Rosa26*^CreERT2/CreERT2^:*Scx*^flox/flox^ mice were treated daily with an intraperitoneal injection of 2mg of tamoxifen (T5648, Millipore Sigma, St. Louis, MO, USA) dissolved in 50µl of corn oil to activate CreERT2 recombinase and generate *Scx*^Δ^ mice (26). *Scx*^+^ mice underwent the same tamoxifen treatment regime, but as they did not have loxP sites flanking scleraxis, no inactivation of scleraxis occurred. Tamoxifen treatment began 5 days prior to synergist ablation procedure or isolation of tendons for cell culture, and continued on a daily basis until tissue was harvested for analysis.

### Synergist ablation tendon growth procedure

Bilateral synergist ablation procedures were performed as described (14, 26, 42). An overview of the time points and surgical procedures are shown in Figure 1B-C. In this procedure, the Achilles tendon is surgically excised. This prevents the gastrocnemius and soleus muscles from plantarflexing the talocrural joint, resulting in compensatory hypertrophy of the adjacent plantaris muscle and tendon. The tendon growth pattern is eccentric, with new tendon tissue, or neotendon, expanding in a mostly lateral direction (Figures 1C and 2A).

Mice were deeply anesthetized with isoflurane, and the hindlimbs were shaved and scrubbed with chlorhexidine solution. A small incision was created in the skin above the posterior hindpaw plantarflexor tendons, and a 3-4mm portion of the Achilles tendon was isolated and excised, leaving remnant stumps at the myotendinous junction and calcaneus. Care was taken to not injure the plantaris tendon during this procedure. The skin was closed with GLUture (Zoetis, Kalamazoo, MI, USA), buprenorphine (0.1mg/kg, subcutaneous) was administered for post-operative analgesia, and *ad libitum* weight-bearing and cage activity were allowed in the postoperative period.

Animals recovered for a period of 7D or 14D, during which time they received daily treatment with tamoxifen. At the end of 7D or 14D, mice were euthanized by exposure to CO2 followed by cervical dislocation. Plantaris tendons were removed and prepared for either DNA isolation, histology, electron microscopy, or RNA isolation. Up to 20 mice per time point and genotype were used in studies.

### Scleraxis Knockdown Efficiency

To determine the efficiency of scleraxis knockdown, DNA was isolated from plantaris tendons or cultured cells using a DNeasy kit (Qiagen, Valencia, CA, USA). DNA was quantified with a NanoDrop 2000 system (Thermo Fisher, Waltham, MA, USA), and qPCR was conducted in a real time thermal cycler using iTaq Universal SYBR Green Supermix (Bio-Rad, Hercules, CA, USA) and primers specific for the region of exon 1 of scleraxis flanked by loxP sites (forward 5’-GACCGCAAGCTCTCCAAGAT-3’; reverse 5’-ACGACCGCTGTGGAAGAAAG-3’), or exon 2 of scleraxis which is outside of the loxP sites (forward 5’-CGCAGGTCCCCAAGAGCACG-3’; reverse 5’-GGCCTGGGTCAGTGTTCGGC-3’). The abundance of exon 1 of scleraxis was measured relative to exon 2 using the 2^-ΔCt^ method, and further normalized to the *Scx*^+^ group for whole tendons, or to *Scx*^+^ tenocytes.

### Histology

Distal portions of plantaris tendons were snap frozen in liquid nitrogen cooled isopentane and stored at -80ºC until use. Tendons were sectioned and stained with fast green and hematoxylin as described previously (14). Fast Green and hematoxylin sections were imaged on a BX51 microscope (Olympus, Center Valley, PA). For immunohistochemistry, tendons were sectioned and permeabilized in Triton-X-100 (0.2%) and blocked with 5% goat serum. Sections were incubated overnight in primary antibodies against CD146 (ab75769, AbCam, Cambridge, MA, USA) and then incubated in goat secondary antibodies conjugated to Alexa Fluor 555 (A32732, Thermo Fisher). Samples were also stained with DAPI (Millipore Sigma) to visualize nuclei and wheat germ agglutinin conjugated to Alexa Fluor 488 (WGA-AF488, W11261, Thermo Fisher) to visualize the ECM. Images were taken on a A1 confocal microscope equipped with a high-resolution camera (Nikon, Tokyo, JP). ImageJ (NIH, Bethesda, MD, USA) was used for quantitative measurements. Cell density was calculated by the number of nuclei per unit area of the tendon. The percentage of CD146^+^ pericyte cells in the neotendon was measured by dividing the total number of CD146^+^/DAPI^+^ cells by the total number of DAPI^+^ cells.

### Transmission Electron Microscopy

Proximal tendon portions were fixed in 1% tannic acid and 1% glutaraldehyde in Sorenson’s buffer, followed by post-fixation in 2% osmium tetroxide (Electron Microscopy Sciences, Hatfield, PA, USA). Samples were then dehydrated using a graded ethanol series and embedded in EMBed 812 resin (Electron Microscopy Sciences) using a graded resin and propylene oxide series. One-micron transverse sections were cut with a diamond knife ultramicrotome, and imaged using a 1400-plus transmission electron microscope with a high-resolution digital camera (JEOL, Peabody, MA, USA). Collagen fibril areas were calculated using ImageJ as described (43).

### Proteomics

Label-free proteomics was performed as reported in previous studies (36, 44) at the University of Liverpool Centre of Genomic Research. Soluble proteins were extracted by homogenizing plantaris tendons in 25mM ammonium bicarbonate and 10mM iodoacetamide, followed by centrifugation and removal of the supernatant. Then 50µg of soluble protein was further reduced and alkylated, and in solution trypsin digestion was performed. The remaining pellet was resuspended in 500µl of 4M Guanidine-HCl extraction buffer (GnHCl), 65mM dithiothreitol, 50mM sodium acetate, pH 5.8 for 48 h at 4ºC with shaking. The samples were then centrifuged, the supernatant was removed, and subjected to in solution-trypsin digest on 10µl of resin (StrataClean, Agilent, Cheshire, UK) followed by reduction and alkylation. The digests (2µL, corresponding to 2.5µg of peptides) were loaded onto a spectrometer (Q-Exactive Quadrupole-Orbitrap, Thermo Scientific) on a one-hour gradient with an intersample 30-minute blank loaded.

Raw spectra were converted to mgf files using Proteome Discovery (Thermo Fisher) and resulting files were searched against the UniProt mouse sequence database using a Mascot server (Matrix Science, London, UK). Search parameters used were: peptide mass tolerance, 10ppm; fragment mass tolerance, 0.01Da; +1,+2,+3 ions; unique peptides ≥2; missed cleavages 1; instrument type ESI-TRAP. Variable modifications included carbamidomethylation of cysteine residues and oxidation of methionine. Label-free quantification was performed using ProgenesisQI software (Waters, Elstree Hertfordshire, UK) as described previously (36, 44). In brief, the MS/MS peak list was searched against the UniProt mouse reviewed database on Mascot using the same search parameters as mentioned above. To account for changes in tendons size, relative protein abundance was normalized to tendon mass, and MetaboAnalyst 4.0 (45) was used to quantify differences between proteins from *Scx*^+^ and *Scx*^Δ^ mice. Proteomics data for this study has been deposited at the ProteomeXchange Consortium via the PRIDE partner repository (accession PXD017944).

### RNA Sequencing and Gene Expression

RNA was isolated from tendons and cultured tenocytes, and quantified as previously described (26, 46). Plantaris tendons or cultured cells were homogenized in QIAzol (Qiagen) and RNA was purified with a miRNeasy Micro Kit (Qiagen) supplemented with DNase I (Qiagen). RNA concentration and quality was determined using a NanoDrop 2000 (Thermo Fisher) and a TapeStation 2200 (Agilent Technologies, Santa Clara, CA, USA). RNA integrity numbers for whole tendon samples were >7.5, and for cultured cells were >9.5.

To sequence RNA, sample concentrations were normalized and cDNA pools were created for each sample, and then subsequently tagged with a barcoded oligo adapter to allow for sample specific resolution. RNAseq of whole tendons was conducted by the University of Michigan Sequencing Core using HiSeq 2500 system (Illumina, San Diego, CA, USA) with 50bp single end reads. RNAseq of cultured cells was conducted by the Weill Cornell Medical College Genomics Core using a NovaSeq 6000 system (Illumina, San Diego, CA, USA) with 50bp paired end reads. Standard Illumina reagents were used for library preparation and sequencing. Raw RNAseq data was quality checked using FastQC v0.10.0 (Barbraham Bioinformatics, Cambridge, UK). Alignment to the reference genome (mmu10, UCSC) was conducted using the STAR aligner, and differential expression was calculated using DESeq2 (47). A false discovery rate (FDR) procedure was applied to correct for multiple observations. Sequencing data has been deposited to NIH GEO (accession GSE145864). Ingenuity Pathway Analysis (IPA) software (Qiagen) was used to perform gene enrichment analysis of RNAseq data described (48, 49).

For qPCR, RNA was reverse transcribed into cDNA using iScript Reverse Transcription reagents (Bio-Rad), and then amplified in a real-time thermal cycler using iTaq Universal SYBR Green Supermix (Bio-Rad). Primer sequences for specific have been previously published (26, 27). Relative copy number of transcripts in the 7D and 14D *Scx*^+^ and *Scx*^Δ^ mice was normalized to 0D non-overloaded *Scx*^+^ controls using the linear regression of efficiency method (50) and log2-transformed prior to analysis.

### Cell culture

Tail tendons were isolated from *Scx*^+^, *Scx*^Δ^, or *Scx*GFP mice, and cells were cultured as described previously (26, 27). Tendon fascicles were carefully isolated from tail tendons, minced, and placed in DMEM with 0.2% type II collagenase (Thermo Scientific) for 1h at 37°C with agitation. CD146^+^ pericytes were isolated from digested cell suspensions of *Scx*^+^ or *Scx*GFP mice using anti-CD146 magnetic microbeads (130-092-007, Miltenyi Biotec, San Diego, CA, USA), and placed into 24-well tissue culture dishes (Corning, Corning, NY, USA). Pericytes were expanded in DMEM containing 10% fetal bovine serum (FBS) and 1% antibiotic-antimycotic (AbAm, Thermo Scientific), referred to as growth medium, for a period of up to 5 days. When experiments involved the evaluation of scleraxis inactivation, growth medium was supplemented with 10μM of 4HT to ensure efficient deletion of *Scx*.

Pericytes from *Scx*GFP mice were placed either directly onto plastic, a thin gel of growth factor-reduced Matrigel (Corning), or type I collagen reconstituted from rat tail tendons as described (27). General cell morphology and GFP abundance from cells of *Scx*GFP mice was measured by visualizing cells in an EVOS FL microscope (Thermo Fisher), and overlaying GFP fluorescence filters with phase contrast images. Pericytes from *Scx*^+^ and *Scx*^Δ^ mice were isolated and cultured on type I collagen gels, and were expanded in growth medium containing 10μM of 4HT. RNA was isolated and gene expression analysis was performed as described above.

Tenocytes were cultured from digested tail tendons, plated on 100mm type I collagen coated dishes (Corning) and expanded in growth medium. Fibroblasts were passaged upon reaching 70% confluence onto 6-well dishes coated with type I collagen (Corning), and media was switched to DMEM containing 2% horse serum and 1% AbAm for a period of two days. RNA was then isolated, and RNAseq and gene expression analysis was performed as described above.

To measure cell proliferation of *Scx*^+^ and *Scx*^Δ^ tenocytes at 50% confluence, 20µM of BrdU (Millipore Sigma) was added to growth media for 1h. Tenocytes were then fixed in ice-cold methanol, permeabilized with 0.5% Triton X-100, and the BrdU epitope was exposed by denaturing DNA using 2 M HCl. Plates were then incubated with anti-BrdU antibodies (G3G4, Developmental Studies Hybridoma Bank, Iowa City, IA, USA) and secondary antibodies conjugated to AF555 (A21127, Thermo Fisher), and DAPI. The number of BrdU^+^ nuclei as a percentage of total nuclei was quantified in 5 randomly selected fields of a single experiment from 6 independent experiments performed per group. Plates were imaged in an EVOS FL microscope and quantified using ImageJ.

### Statistics

Results are presented as mean±standard deviation (SD) or mean±coefficient of variation (CV). Prism version 8.0 (GraphPad Software, La Jolla, CA, USA) was used to conduct statistical analyses for all data except RNAseq. A two-way ANOVA followed by Fisher’s *post hoc* sorting (α=0.05) evaluated the interaction between time after synergist ablation and *Scx* knockdown. For cell culture experiments, differences between groups were tested with an unpaired Student’s t-test (α=0.05), or a one- or two-way ANOVA followed by Fisher’s *post hoc* sorting (α=0.05). A Pearson correlation coefficient (r) was calculated for the percentage of CD146^+^ cells in the neotendon as a function of neotendon CSA. For CSA quantification of fibrils, differences between the bins of the two groups were tested with a Chi-squared test (α=0.05). Replicates for whole tissue experiments are tendons from individual animals, while cell culture experiments represent technical replicates of individual or pooled animals.

### Study Approval

Animal studies were approved by the University of Michigan (protocol PRO00003566) and Hospital for Special Surgery/Weill Cornell Medical College/Memorial Sloan Kettering (protocol 2017-0035) Institutional Animal Care & Use Committees. Experimental procedures were conducted according to approved protocols and in accordance with the US Public Health Policies on Human Care and Use of Laboratory Animals.

## Author Contributions

JPG, MMS, and CLM designed research; JPG, MMS, and YAK performed research; JPG, MMS, YAK, EC, and CLM analyzed data; JPG, MMS, and CLM drafted and edited the manuscript; all authors approved of the final version of the manuscript.

## Acknowledgements

The authors would like to acknowledge technical assistance from Dr. Deborah Simpson at the University of Liverpool. The *Scx*GFP and *Scx*^flox^ mice used in this study were kindly provided by Dr. Ronen Schweitzer at the Shriners Hospital for Children in Portland, Oregon. This work was supported by grants R01-AR063649 and F31-AR065931 from the National Institutes of Health.

