## Supplemental Material for "Scleraxis is required for the growth of adult tendons in response to mechanical loading"

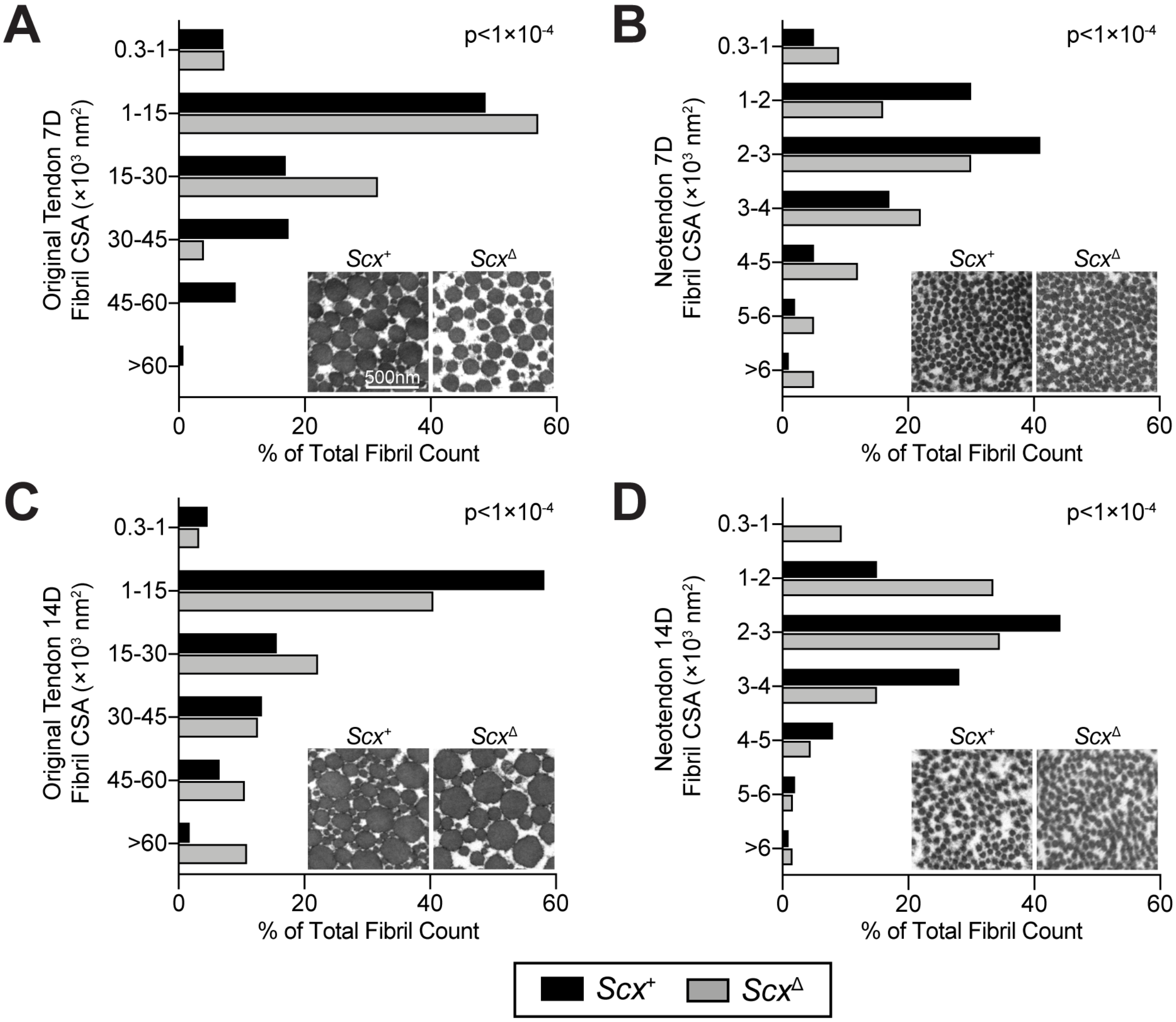


***Supplemental Figure 1. Effect of scleraxis deletion on tendon collagen fibril size distributions.*** Quantification of collagen fibril cross-sectional area (CSA) measurements, from transmission electron micrographs of *Scx*$^{+}$ and *Scx*$^{\Delta}$ (A, C) original tendons or (B, D) neotendons at either (A, B) 7D or (C, D) 14D after synergist ablation/plantaris growth procedure. Fibril counts are expressed as percentage of total fibril abundance. Scale bar for all images is 500nm. Differences between the distribution of *Scx*$^{+}$ and *Scx*$^{\Delta}$ fibrils for each time point and tendon area were tested using the Chi-squared test ($p<0.05$). N≥2 mice per group.

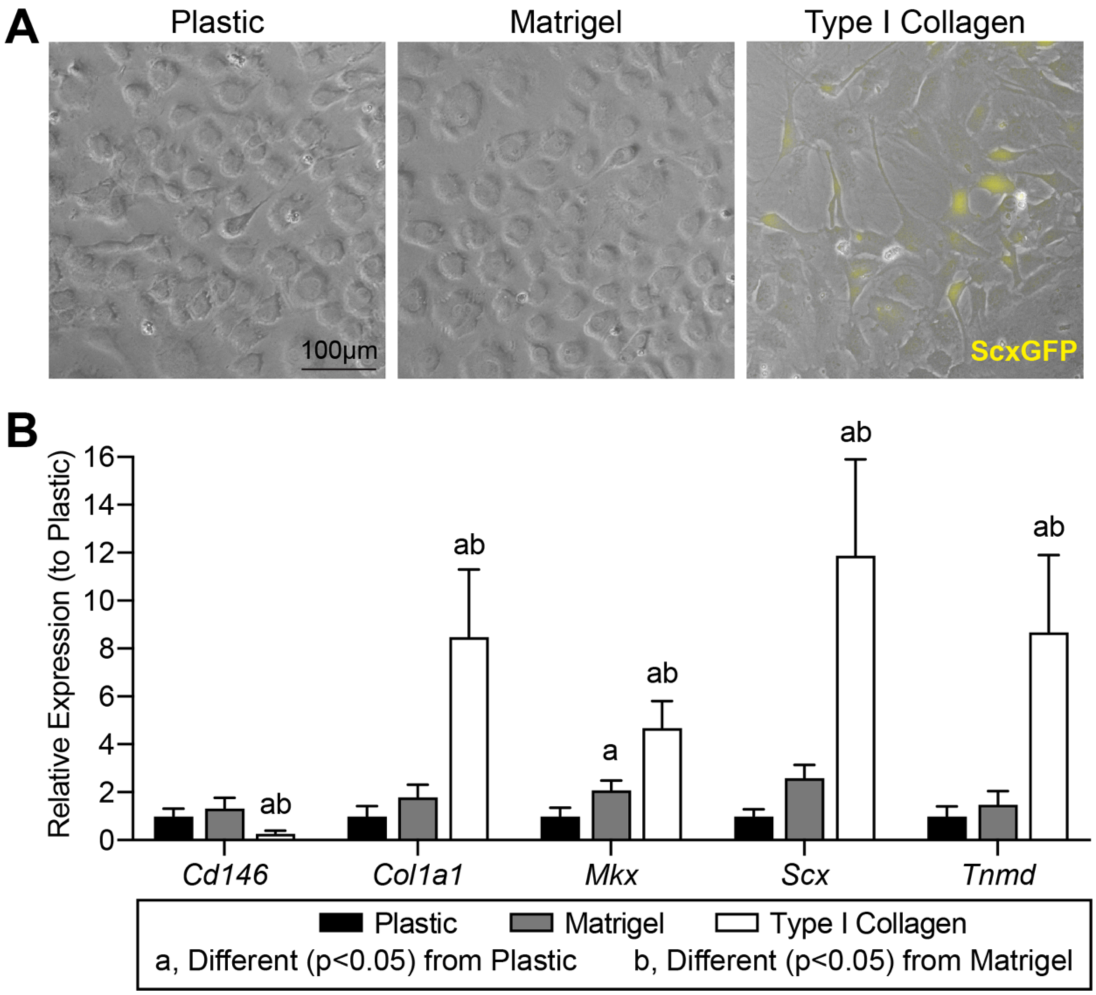


***Supplemental Figure 2. Effect of matrix substrate on pericyte differentiation.*** (A) CD146$^+$ pericytes from *Scx*GFP mice were cultured on plastic, or on thin gels of Matrigel or type I collagen, for 5 days. Fluorescent images demonstrate GFP expression (driven by scleraxis, shown in yellow), with phase contrast overlay. Scale bar for all images is 100µm. (B) Gene expression of pericytes placed on different culture substrates for 5 days. Values are mean±SD. Differences between groups were tested using a two-way ANOVA: a, significantly different ($p<0.05$) from Plastic; b, significantly different ($p<0.05$) from Matrigel. N=6 replicates per group.

***Supplemental Table 1.*** Effect of scleraxis deletion on the proteome of tendons. Abundance of proteins in *Scx*$^{\Delta}$ mice normalized to *Scx*$^{+}$ mice.

| Protein | **7D *Scx*$^{\Delta}$/*Scx*$^{+}$ Log$_2$ fold change** | ***7D Scx*$^{\Delta}$/*Scx*$^{+}$ q value** | **14D *Scx*$^{\Delta}$/*Scx*$^{+}$ Log$_2$ fold change** | ***14D Scx*$^{\Delta}$/*Scx*$^{+}$ q value** |
|---|---|---|---|---|
| Acaa2 | 0.3312 | *>0.999999* | 1.368 | *0.02115* |
| Acads | -0.2764 | *>0.999999* | 1.56 | *0.014388* |
| Acadvl | -0.07899 | *>0.999999* | 1.305 | *0.022838* |
| Acat1 | 0.5886 | *>0.999999* | 0.9696 | *0.05219* |
| Aco2 | 0.07635 | *>0.999999* | 0.7631 | *0.097758* |
| Acta1 | -1.809 | *0.223308* | 1.123 | *0.032381* |
| Actb | -0.4552 | *>0.999999* | 1.344 | *0.021333* |
| Actc1 | -1.953 | *0.18802* | 1.305 | *0.022838* |
| Actn1 | 0.5011 | *>0.999999* | 1.417 | *0.019205* |
| Actn2 | -0.04605 | *>0.999999* | 1.138 | *0.030905* |
| Actn3 | -0.1859 | *>0.999999* | 0.9337 | *0.057667* |
| Actn4 | 0.8351 | *0.978994* | 1.164 | *0.029612* |
| Adssl1 | 0.6943 | *>0.999999* | 0.9351 | *0.057667* |
| Aebp1 | -0.2623 | *>0.999999* | 0.9497 | *0.055263* |
| Ahsg | 0.8818 | *0.962632* | 0.6694 | *0.127609* |
| Ak1 | -0.2752 | *>0.999999* | 1.062 | *0.039606* |
| Alb | 1.396 | *0.547167* | 1.159 | *0.029986* |
| Aldh1b1 | 1.767 | *0.244839* | 1.269 | *0.02361* |
| Aldh2 | 0.5773 | *>0.999999* | 1.124 | *0.032307* |
| Aldoa | -0.4571 | *>0.999999* | 0.9514 | *0.055044* |
| Ampd1 | 1.27 | *0.697595* | -0.3223 | *0.31011* |
| Angptl2 | 0.04023 | *>0.999999* | 0.6883 | *0.120875* |
| Angptl7 | 0.6807 | *>0.999999* | 1.495 | *0.01684* |
| Anxa1 | 0.6343 | *>0.999999* | 0.9868 | *0.049131* |
| Anxa2 | 0.5449 | *>0.999999* | 0.7939 | *0.088909* |
| Anxa5 | 0.3185 | *>0.999999* | 1.358 | *0.021333* |
| Anxa6 | 0.02231 | *>0.999999* | 0.7504 | *0.100486* |
| Ap2a1 | 0.5921 | *>0.999999* | 0.4648 | *0.222708* |
| Apobec2 | -1.988 | *0.212303* | 0.2116 | *0.388929* |
| Apoh | 0.9011 | *0.950388* | 1.051 | *0.041055* |
| Arcn1 | 0.8991 | *0.950388* | 1.498 | *0.016799* |
| Aspn | 0.9297 | *0.936877* | 1.263 | *0.02383* |
| Atp2a1 | -0.4819 | *>0.999999* | 0.5977 | *0.154893* |
| Atp5a1 | 0.03157 | *>0.999999* | 1.037 | *0.042437* |
| Atp5b | 0.1115 | *>0.999999* | 1.039 | *0.042426* |
| Atp5c1 | 0.2423 | *>0.999999* | 1.049 | *0.041055* |
| Atp5h | 0.02411 | *>0.999999* | 1.229 | *0.025112* |
| Atp5i | 0.9434 | *0.928916* | 2.287 | *0.00063* |
| Atp5o | 0.004643 | *>0.999999* | 0.8854 | *0.067416* |
| Bgn | 1.002 | *0.905842* | 0.6258 | *0.143872* |
| Bin1 | -1.293 | *0.663911* | 0.7542 | *0.1004* |
| C1qtnf3 | 0.4504 | *>0.999999* | 1.901 | *0.003928* |
| Ca3 | -0.3194 | *>0.999999* | 0.1173 | *0.464192* |
| Calr | -0.2091 | *>0.999999* | 1.157 | *0.030101* |
| Camk2a | 0.8632 | *0.962632* | 0.8269 | *0.080864* |
| Camp | 1.223 | *0.709595* | -0.31 | *0.317714* |
| Capza2 | -0.5417 | *>0.999999* | 1.102 | *0.034231* |
| Capzb | 0.5401 | *>0.999999* | 0.8754 | *0.069052* |
| Casq1 | -1.45 | *0.505056* | 1.242 | *0.024265* |
| Cbr2 | 0.202 | *>0.999999* | 1.139 | *0.030854* |
| Cct2 | -0.6017 | *>0.999999* | 1.651 | *0.011346* |
| Cct3 | 0.1214 | *>0.999999* | 1.442 | *0.01879* |
| Cct4 | -0.08833 | *>0.999999* | 1.65 | *0.011346* |
| Cct5 | 0.2762 | *>0.999999* | 1.476 | *0.018026* |
| Cct8 | -0.1507 | *>0.999999* | 1.534 | *0.015815* |
| Cd36 | 0.589 | *>0.999999* | 1.311 | *0.022838* |
| Cfl1 | -0.1991 | *>0.999999* | 1.678 | *0.010958* |
| Cfl2 | 0.4622 | *>0.999999* | 2.117 | *0.003025* |
| Chad | 2.783 | *0.01017* | 1.983 | *0.002731* |
| Cilp | 3.499 | *0.00034* | 4.81 | *<0.000001* |

| | | | | |
|---|---|---|---|---|
| Ckap4 | 0.9376 | *0.928916* | 1.703 | *0.009895* |
| Ckm | -1.402 | *0.547167* | -0.4127 | *0.252462* |
| Ckmt2 | -0.7881 | *>0.999999* | 0.9709 | *0.05219* |
| Cltc | -0.2257 | *>0.999999* | 0.5835 | *0.161131* |
| Clu | -0.9467 | *0.928916* | 1.288 | *0.022838* |
| Col11a1 | 1.227 | *0.709595* | 1.77 | *0.008253* |
| Col12a1 | 0.4596 | *>0.999999* | 1.05 | *0.041055* |
| Col14a1 | -0.08529 | *>0.999999* | 0.278 | *0.340116* |
| Col1a1 | 0.6588 | *>0.999999* | 0.6159 | *0.147619* |
| Col1a2 | 0.666 | *>0.999999* | 0.6141 | *0.147774* |
| Col2a1 | 0.6961 | *>0.999999* | 0.6504 | *0.134584* |
| Col3a1 | 1.843 | *0.212303* | 1.021 | *0.044597* |
| Col5a1 | 1.483 | *0.483526* | 1.645 | *0.011416* |
| Col5a2 | 1.375 | *0.562088* | 1.201 | *0.026427* |
| Col6a1 | 0.7241 | *>0.999999* | 0.9302 | *0.057989* |
| Col6a2 | 0.8611 | *0.962632* | 0.8648 | *0.071132* |
| Colgalt1 | -0.08009 | *>0.999999* | 2.195 | *0.001082* |
| Comp | 0.8566 | *0.962632* | 0.8798 | *0.068426* |
| Copa | 0.1625 | *>0.999999* | 1.254 | *0.02392* |
| Coro1c | 0.7284 | *>0.999999* | 1.368 | *0.02115* |
| Cox4i1 | -0.1971 | *>0.999999* | 1.067 | *0.039027* |
| Cox6c | 0.7238 | *>0.999999* | 0.9541 | *0.054754* |
| Cryab | -2.829 | *0.008995* | 0.6378 | *0.138882* |
| Cs | -0.4499 | *>0.999999* | 1.295 | *0.022838* |
| Csrp3 | -1.928 | *0.190099* | -0.8201 | *0.082485* |
| Ctsk | 0.7666 | *>0.999999* | 1.112 | *0.03321* |
| Cyc1 | -0.159 | *>0.999999* | 1.139 | *0.030854* |
| Cycs | 0.05033 | *>0.999999* | 1.464 | *0.018026* |
| D1Pas1 | 0.1121 | *>0.999999* | 1.507 | *0.016633* |
| Dcn | 1.145 | *0.791754* | 0.884 | *0.06762* |
| Ddost | 0.3134 | *>0.999999* | 1.171 | *0.029043* |
| Ddx1 | 0.3734 | *>0.999999* | 1.258 | *0.02392* |
| Ddx17 | -1.514 | *0.462906* | 1.804 | *0.007433* |
| Ddx5 | -0.2238 | *>0.999999* | 1.317 | *0.022655* |
| Des | -1.519 | *0.462906* | 1.193 | *0.027057* |
| Dlat | -0.5864 | *>0.999999* | 1.083 | *0.036523* |
| Dld | 1.097 | *0.818857* | 1.016 | *0.044597* |
| Dlst | 0.158 | *>0.999999* | 1.267 | *0.023619* |
| Dpt | 0.3796 | *>0.999999* | 0.8026 | *0.087007* |
| Dpysl2 | 0.6188 | *>0.999999* | 1.277 | *0.023131* |
| Dpysl3 | -0.1346 | *>0.999999* | 1.058 | *0.039836* |
| Eef1a1 | 0.1351 | *>0.999999* | 1.74 | *0.009308* |
| Eef1a2 | -0.6437 | *>0.999999* | -0.3013 | *0.323372* |
| Eef1g | -0.8247 | *0.978994* | 1.712 | *0.009647* |
| Eef2 | 0.3378 | *>0.999999* | 1.798 | *0.007433* |
| Ehd2 | -0.449 | *>0.999999* | 1.32 | *0.022443* |
| Ehd4 | -0.8544 | *0.962632* | 1.262 | *0.02383* |
| Eif4a1 | 0.6565 | *>0.999999* | 1.526 | *0.016246* |
| Emilin1 | 0.1746 | *>0.999999* | 0.8469 | *0.075258* |
| Eno1 | -0.1141 | *>0.999999* | 1.396 | *0.019847* |
| Eno2 | -2.572 | *0.04782* | -0.9487 | *0.055337* |
| Eno3 | -0.9919 | *0.905842* | -0.4543 | *0.227609* |
| Eprs | 0.007425 | *>0.999999* | 1.187 | *0.027445* |
| Esd | -0.2585 | *>0.999999* | 1.227 | *0.025112* |
| Etfa | 0.2896 | *>0.999999* | 1.161 | *0.029979* |
| Etfb | 0.2333 | *>0.999999* | 1.285 | *0.022838* |
| F13a1 | 0.2307 | *>0.999999* | 0.7099 | *0.113346* |
| Fbl | 0.1449 | *>0.999999* | 1 | *0.047131* |
| Fbn1 | -0.6105 | *>0.999999* | 1.105 | *0.033945* |
| Fbn2 | 0.1034 | *>0.999999* | 0.8005 | *0.087409* |
| Fga | 0.7101 | *>0.999999* | 1.194 | *0.027017* |
| Fgb | 1.003 | *0.905842* | 1.235 | *0.024785* |
| Fgg | 0.9135 | *0.950388* | 1.349 | *0.021333* |
| Fh | 0.5207 | *>0.999999* | 1.075 | *0.037803* |
| Fhl1 | -1.641 | *0.351025* | 0.3356 | *0.301167* |
| Fkbp10 | 0.3989 | *>0.999999* | 1.439 | *0.01879* |

| | | | | |
|---|---|---|---|---|
| Flna | 0.872 | 0.962632 | 1.337 | 0.021629 |
| Flnb | 0.2289 | >0.999999 | 1.343 | 0.021333 |
| Flnc | -1.511 | 0.462906 | 1.003 | 0.046756 |
| Fmod | 0.8247 | 0.978994 | 0.9213 | 0.059446 |
| Fn1 | 0.1989 | >0.999999 | 1.686 | 0.010712 |
| Gapdh | 0.2193 | >0.999999 | 0.9888 | 0.048855 |
| Glud1 | 0.9663 | 0.921594 | 0.5426 | 0.181605 |
| Gnb1 | 0.7584 | >0.999999 | 0.8901 | 0.066553 |
| Gnb2 | 0.07633 | >0.999999 | 0.933 | 0.057667 |
| Gnb2l1 | 0.3735 | >0.999999 | 1.242 | 0.024265 |
| Got2 | 0.319 | >0.999999 | 1.27 | 0.02361 |
| Gpd1 | 2.668 | 0.036698 | -1.325 | 0.025112 |
| Gpd2 | -0.3109 | >0.999999 | 1.232 | 0.0249 |
| Gsn | 0.08789 | >0.999999 | 1.023 | 0.044403 |
| H1f0 | 0.29 | >0.999999 | 0.7976 | 0.087915 |
| H2afy | 0.5526 | >0.999999 | 0.8733 | 0.069354 |
| H2afz | 0.9145 | 0.950388 | 0.7614 | 0.098127 |
| H3f3c | -2.88 | 0.007804 | -0.9705 | 0.05219 |
| Hadha | 0.08571 | >0.999999 | 1.343 | 0.021333 |
| Hadhb | -0.03645 | >0.999999 | 0.9683 | 0.05219 |
| Hba | -0.05102 | >0.999999 | 0.751 | 0.100486 |
| Hbb-b1 | -0.2273 | >0.999999 | 0.7516 | 0.100486 |
| Hdlbp | 0.09162 | >0.999999 | 1.515 | 0.016633 |
| Hist1h1a | -1.101 | 0.818857 | 0.8627 | 0.07151 |
| Hist1h1b | -0.8592 | 0.962632 | 1.037 | 0.042437 |
| Hist1h1c | -0.7377 | >0.999999 | 0.7842 | 0.091209 |
| Hist1h1e | -0.3852 | >0.999999 | 0.7999 | 0.087409 |
| Hist1h2ab | -0.5032 | >0.999999 | 0.999 | 0.047202 |
| Hist1h2bc | 0.7051 | >0.999999 | 0.7769 | 0.093424 |
| Hist1h3b | -1.671 | 0.325362 | -1.198 | 0.026621 |
| Hist1h4a | 0.6279 | >0.999999 | 0.8168 | 0.083101 |
| Hist2h2aa1 | 0.4164 | >0.999999 | 1.017 | 0.044597 |
| Hist2h2be | 4.877 | <0.000001 | 1.322 | 0.022395 |
| Hnrnpa0 | -0.9935 | 0.905842 | 1.564 | 0.014388 |
| Hnrnpa1 | 0.2597 | >0.999999 | 0.7138 | 0.112101 |
| Hnrnpa3 | 0.1195 | >0.999999 | 1.066 | 0.039027 |
| Hnrnph1 | -0.1115 | >0.999999 | 0.9365 | 0.057527 |
| Hnrnpl | 1.302 | 0.658837 | 0.8161 | 0.083101 |
| Hnrnpm | 0.6691 | >0.999999 | 1.156 | 0.030101 |
| Hnrnpu | -0.2155 | >0.999999 | 1.326 | 0.022106 |
| Hpx | 2.056 | 0.149859 | 1.57 | 0.014388 |
| Hsp90aa1 | -0.3604 | >0.999999 | 1.523 | 0.016286 |
| Hsp90ab1 | -0.3124 | >0.999999 | 1.374 | 0.021037 |
| Hsp90b1 | 0.0738 | >0.999999 | 1.125 | 0.032307 |
| Hspa1b | -0.2878 | >0.999999 | 0.8597 | 0.072178 |
| Hspa5 | 0.8279 | 0.978994 | 1.283 | 0.022838 |
| Hspa8 | 0.06885 | >0.999999 | 1.016 | 0.044597 |
| Hspa9 | 0.5894 | >0.999999 | 0.8676 | 0.070546 |
| Hspb1 | -1.941 | 0.189038 | 0.5159 | 0.195039 |
| Hspb7 | 0.1389 | >0.999999 | 1.358 | 0.021333 |
| Hspd1 | 0.1009 | >0.999999 | 1.248 | 0.024171 |
| Hspg2 | -0.1178 | >0.999999 | 1.282 | 0.022838 |
| Idh1 | 0.2311 | >0.999999 | 1.468 | 0.018026 |
| Idh3a | 0.2629 | >0.999999 | 1.139 | 0.030854 |
| Idh3g | -0.5495 | >0.999999 | 1.172 | 0.029043 |
| Ig gamma-2A chain C region | -1.172 | 0.757543 | 1.098 | 0.034362 |
| Ig kappa chain C region | 0.3746 | >0.999999 | 1.148 | 0.030535 |
| Ighm | 1.402 | 0.547167 | 2.483 | 0.000178 |
| Ikbip | 0.2396 | >0.999999 | 1.501 | 0.016782 |
| Immt | -0.02585 | >0.999999 | 0.934 | 0.057667 |
| Iqgap1 | -0.1597 | >0.999999 | 1.506 | 0.016633 |
| Itih1 | 0.4955 | >0.999999 | 0.6244 | 0.143989 |
| Itih3 | 1.026 | 0.889665 | 0.6095 | 0.149592 |
| Jph1 | 0.3308 | >0.999999 | 1.136 | 0.030983 |
| Jph2 | -0.5969 | >0.999999 | 1.093 | 0.035014 |
| Jup | 1.087 | 0.820252 | -0.6663 | 0.128611 |

| | | | | |
|---|---|---|---|---|
| Kera | 1.167 | *0.757543* | 1.026 | *0.044106* |
| Klhl41 | -0.5146 | *>0.999999* | 1.007 | *0.046196* |
| Krt1 | 1.678 | *0.325362* | -0.4166 | *0.250518* |
| Krt10 | 2.196 | *0.102058* | -0.5399 | *0.182605* |
| Krt16 | 1.826 | *0.216659* | -0.05127 | *0.519199* |
| Krt17 | 2.047 | *0.149859* | -0.217 | *0.385988* |
| Krt2 | 2.996 | *0.004777* | -0.3461 | *0.294978* |
| Krt31 | 0.6943 | *>0.999999* | 0.7466 | *0.101348* |
| Krt32 | -1.466 | *0.499658* | 1.47 | *0.018026* |
| Krt34 | 1.339 | *0.602081* | 0.228 | *0.378096* |
| Krt42 | 1.903 | *0.196129* | -0.5924 | *0.157075* |
| Krt5 | 0.7519 | *>0.999999* | 0.4549 | *0.227609* |
| Krt6a | 0.1362 | *>0.999999* | 0.896 | *0.065178* |
| Krt73 | 0.1663 | *>0.999999* | -0.9931 | *0.048111* |
| Krt76 | 2.027 | *0.154328* | -0.4281 | *0.243594* |
| Krt79 | 2.441 | *0.038908* | -0.3108 | *0.317714* |
| Krt82 | -1.34 | *0.602081* | 0.7039 | *0.115431* |
| Krt85 | -0.5969 | *>0.999999* | 0.7186 | *0.11079* |
| Krt86 | 1.211 | *0.722907* | 2.363 | *0.000404* |
| Lama2 | 0.1038 | *>0.999999* | 1.291 | *0.022838* |
| Ldb3 | -0.5482 | *>0.999999* | 1.188 | *0.027445* |
| Ldha | 0.4229 | *>0.999999* | 1.336 | *0.021629* |
| Lima1 | -0.1224 | *>0.999999* | 1.437 | *0.01879* |
| Lmna | 0.1506 | *>0.999999* | 0.6389 | *0.138724* |
| Lmnb1 | 0.3812 | *>0.999999* | 0.8195 | *0.082485* |
| Lox | 1.185 | *0.754322* | 1.652 | *0.011346* |
| Lrp1 | 0.7537 | *>0.999999* | 0.6615 | *0.130352* |
| Ltf | 2.51 | *0.033567* | -0.4721 | *0.218776* |
| Lum | 0.5619 | *>0.999999* | 0.4638 | *0.222803* |
| Macrod1 | -0.9377 | *0.928916* | 2.075 | *0.002067* |
| Matn2 | 0.2389 | *>0.999999* | 0.5022 | *0.202406* |
| Mdh2 | 0.5545 | *>0.999999* | 1.21 | *0.026089* |
| Mfge8 | 1.229 | *0.709595* | 1.434 | *0.018948* |
| Mmp2 | 0.5521 | *>0.999999* | 1.147 | *0.030624* |
| Mpo | 0.3096 | *>0.999999* | 0.9942 | *0.048043* |
| Msn | 0.3695 | *>0.999999* | 1.143 | *0.030833* |
| Mtco2 | 0.08718 | *>0.999999* | 1.505 | *0.016633* |
| Mvp | 0.4298 | *>0.999999* | 1.116 | *0.033085* |
| Mybpc2 | -0.7974 | *0.998346* | 1.018 | *0.044597* |
| Myh1 | -0.9906 | *0.905842* | 0.9706 | *0.05219* |
| Myh10 | -0.9381 | *0.928916* | 1.1 | *0.034288* |
| Myh3 | -1.775 | *0.244839* | 1.289 | *0.022838* |
| Myh4 | -1.894 | *0.196129* | 0.905 | *0.063195* |
| Myh8 | -0.4164 | *>0.999999* | 1.234 | *0.024785* |
| Myh9 | 0.4202 | *>0.999999* | 1.269 | *0.02361* |
| Myl1 | -2.05 | *0.149859* | 1.117 | *0.033085* |
| Myl12b | 0.1057 | *>0.999999* | 1.351 | *0.021333* |
| Myl4 | -3.443 | *0.00039* | 1.227 | *0.030983* |
| Myl6 | 1.065 | *0.843552* | 1.024 | *0.044339* |
| Myl6b | -4.435 | *0.000045* | 3.899 | *<0.000001* |
| Mylpf | 0.003785 | *>0.999999* | 0.6515 | *0.13443* |
| Myo1c | 0.3335 | *>0.999999* | 0.9453 | *0.055946* |
| Myoc | -0.8095 | *0.995305* | -0.2002 | *0.396845* |
| Myom1 | -0.05239 | *>0.999999* | 0.7143 | *0.112101* |
| Myom3 | 0.2785 | *>0.999999* | 0.537 | *0.183733* |
| Myot | -0.9668 | *0.921594* | 0.7471 | *0.101348* |
| Myoz1 | -0.05961 | *>0.999999* | 0.7762 | *0.093427* |
| Naca | 0.9482 | *0.928916* | 0.4115 | *0.252671* |
| Ncl | 0.7297 | *>0.999999* | 0.9088 | *0.062405* |
| Ndufa10 | -0.3214 | *>0.999999* | 1.115 | *0.033151* |
| Ndufa13 | -0.2805 | *>0.999999* | 0.9271 | *0.058559* |
| Ndufa4 | -0.1943 | *>0.999999* | 1.255 | *0.02392* |
| Ndufa9 | 0.4618 | *>0.999999* | 1.927 | *0.003431* |
| Ndufb10 | -0.193 | *>0.999999* | 0.6145 | *0.147774* |
| Ndufs1 | -0.5817 | *>0.999999* | 1.017 | *0.044597* |
| Ndufs3 | -0.5128 | *>0.999999* | 1.312 | *0.022838* |

| | | | | |
|---|---|---|---|---|
| Ndufv1 | -0.04459 | >0.999999 | 1.038 | 0.042426 |
| Ngp | 5.535 | <0.000001 | -1.13 | 0.031767 |
| Nid1 | 0.7785 | >0.999999 | 0.8081 | 0.085426 |
| Npm1 | 1.541 | 0.452588 | 1.442 | 0.01879 |
| Nucb1 | 1.5 | 0.469473 | 1.394 | 0.019857 |
| Nucb2 | 2.078 | 0.149859 | 1.586 | 0.014301 |
| Oat | 0.4092 | >0.999999 | 1.409 | 0.019503 |
| Obscn | -0.5688 | >0.999999 | 0.6919 | 0.11972 |
| Ogdh | 0.4878 | >0.999999 | 1.439 | 0.01879 |
| Ogn | 0.7384 | >0.999999 | 1.04 | 0.042395 |
| P4ha1 | 1.119 | 0.818857 | 1.598 | 0.013906 |
| P4hb | 1.427 | 0.528353 | 1.172 | 0.029043 |
| Pa2g4 | -2.507 | 0.033567 | 1.286 | 0.022838 |
| Pabpc1 | 0.6349 | >0.999999 | 0.9553 | 0.054662 |
| Park7 | 0.7837 | >0.999999 | 0.9303 | 0.057989 |
| Pbxip1 | -0.8643 | 0.962632 | 0.8573 | 0.072452 |
| Pcolce | 1.238 | 0.709595 | 0.9001 | 0.064295 |
| Pdha1 | 0.4118 | >0.999999 | 0.8796 | 0.068426 |
| Pdhb | 0.03031 | >0.999999 | 1.2 | 0.026481 |
| Pdia3 | 0.428 | >0.999999 | 1.148 | 0.030535 |
| Pdia6 | 0.02544 | >0.999999 | 1.182 | 0.027999 |
| Pdlim3 | -0.0162 | >0.999999 | 0.3833 | 0.269734 |
| Pdlim5 | -0.8993 | 0.950388 | 1.061 | 0.039648 |
| Pdlim7 | -0.3776 | >0.999999 | 1.728 | 0.009403 |
| Pfkm | -0.71 | >0.999999 | 1.1 | 0.034288 |
| Pfn1 | 1.03 | 0.889665 | 1.332 | 0.021762 |
| Pgam2 | 1.061 | 0.843552 | 0.8757 | 0.069052 |
| Pgk1 | 0.9818 | 0.906848 | 0.9668 | 0.052373 |
| Phb | 0.6607 | >0.999999 | 0.9256 | 0.058762 |
| Phb2 | 0.3923 | >0.999999 | 0.886 | 0.067416 |
| Pkm | 0.3175 | >0.999999 | 1.462 | 0.018026 |
| Plec | 0.3125 | >0.999999 | 0.7423 | 0.10268 |
| Postn | -0.5579 | >0.999999 | 1.555 | 0.014388 |
| Ppia | 1.076 | 0.832698 | 1.006 | 0.046284 |
| Ppib | 0.8207 | 0.979766 | 1.405 | 0.019503 |
| Prdx1 | 0.6731 | >0.999999 | 1.39 | 0.019857 |
| Prdx2 | 1.224 | 0.709595 | 1.208 | 0.026089 |
| Prelp | 0.8523 | 0.962632 | 0.3402 | 0.298507 |
| Prg4 | 0.1705 | >0.999999 | 0.9386 | 0.057371 |
| Prkcdbp | -0.1624 | >0.999999 | 0.731 | 0.106408 |
| Psmc3 | -0.01791 | >0.999999 | 1.219 | 0.025578 |
| Psmd12 | -0.8781 | 0.962632 | 2.184 | 0.001082 |
| Ptbp1 | 0.4184 | >0.999999 | 1.508 | 0.016633 |
| Ptrf | -0.9956 | 0.905842 | 0.7534 | 0.1004 |
| Pygm | 0.1921 | >0.999999 | 0.7529 | 0.1004 |
| Rab10 | 0.5073 | >0.999999 | 1.633 | 0.012009 |
| Rab14 | 0.9867 | 0.905842 | 1.102 | 0.034231 |
| Rab1A | -0.1018 | >0.999999 | 1.558 | 0.014388 |
| Rab7a | 0.3504 | >0.999999 | 1.21 | 0.026089 |
| Ran | 0.6006 | >0.999999 | 1.228 | 0.025112 |
| Rap1b | 0.1778 | >0.999999 | 1.309 | 0.022838 |
| Rhoa | 1.17 | 0.757543 | 1.165 | 0.029612 |
| Rhoc | -0.7207 | >0.999999 | 1.655 | 0.011346 |
| Rpl10 | -0.09035 | >0.999999 | 1.663 | 0.011346 |
| Rpl10a | 0.4289 | >0.999999 | 1.416 | 0.019205 |
| Rpl11 | -0.4812 | >0.999999 | 1.254 | 0.02392 |
| Rpl12 | -0.3488 | >0.999999 | 1.3 | 0.022838 |
| Rpl13 | -0.006 | >0.999999 | 1.307 | 0.022838 |
| Rpl13a | 0.1437 | >0.999999 | 1.188 | 0.027445 |
| Rpl14 | 0.4214 | >0.999999 | 1.267 | 0.023619 |
| Rpl15 | 0.4179 | >0.999999 | 1.326 | 0.022106 |
| Rpl17 | -0.00462 | >0.999999 | 1.351 | 0.021333 |
| Rpl18 | 1.035 | 0.889665 | 0.8383 | 0.077599 |
| Rpl18a | 0.678 | >0.999999 | 1.276 | 0.023187 |
| Rpl19 | -0.00961 | >0.999999 | 1.26 | 0.02383 |
| Rpl21 | 0.4268 | >0.999999 | 1.141 | 0.030854 |

| | | | | |
|---|---|---|---|---|
| Rpl22 | 0.3005 | *>0.999999* | 1.201 | *0.026427* |
| Rpl23 | 0.1949 | *>0.999999* | 1.365 | *0.021325* |
| Rpl23a | -0.2316 | *>0.999999* | 1.466 | *0.018026* |
| Rpl24 | 0.01345 | *>0.999999* | 1.422 | *0.019205* |
| Rpl26 | -0.07845 | *>0.999999* | 1.214 | *0.026042* |
| Rpl27 | 0.006702 | *>0.999999* | 1.423 | *0.019205* |
| Rpl27a | 0.06541 | *>0.999999* | 1.377 | *0.020842* |
| Rpl28 | -0.06622 | *>0.999999* | 1.292 | *0.022838* |
| Rpl3 | 0.1549 | *>0.999999* | 1.354 | *0.021333* |
| Rpl30 | 0.2454 | *>0.999999* | 1.221 | *0.025536* |
| Rpl31 | 0.02629 | *>0.999999* | 1.368 | *0.02115* |
| Rpl32 | -0.4282 | *>0.999999* | 1.244 | *0.024265* |
| Rpl34 | -0.2576 | *>0.999999* | 1.145 | *0.030677* |
| Rpl35 | -0.3059 | *>0.999999* | 1.611 | *0.013473* |
| Rpl35a | 0.4301 | *>0.999999* | 1.224 | *0.025298* |
| Rpl36 | -0.6446 | *>0.999999* | 1.578 | *0.014379* |
| Rpl36a | 0.1183 | *>0.999999* | 1.282 | *0.022838* |
| Rpl37a | -0.4227 | *>0.999999* | 1.422 | *0.019205* |
| Rpl38 | 0.07964 | *>0.999999* | 1.443 | *0.01879* |
| Rpl4 | 0.4061 | *>0.999999* | 1.294 | *0.022838* |
| Rpl5 | 0.2597 | *>0.999999* | 1.316 | *0.022666* |
| Rpl6 | 0.5123 | *>0.999999* | 1.405 | *0.019503* |
| Rpl7 | 0.2888 | *>0.999999* | 0.9689 | *0.05219* |
| Rpl7a | -0.05992 | *>0.999999* | 1.294 | *0.022838* |
| Rpl8 | 0.2876 | *>0.999999* | 1.3 | *0.022838* |
| Rpl9 | -0.1854 | *>0.999999* | 1.432 | *0.018948* |
| Rplp0 | 0.02288 | *>0.999999* | 1.587 | *0.014301* |
| Rpn1 | 0.8325 | *0.978994* | 1.253 | *0.02392* |
| Rpn2 | 1.55 | *0.450704* | 1.041 | *0.042395* |
| Rps10 | -0.1926 | *>0.999999* | 1.583 | *0.014301* |
| Rps11 | 0.1459 | *>0.999999* | 1.399 | *0.019794* |
| Rps12 | 0.6131 | *>0.999999* | 1.181 | *0.027999* |
| Rps13 | 0.4003 | *>0.999999* | 1.338 | *0.021629* |
| Rps14 | 0.1842 | *>0.999999* | 1.465 | *0.018026* |
| Rps15a | 0.4331 | *>0.999999* | 1.468 | *0.018026* |
| Rps16 | 0.3298 | *>0.999999* | 1.21 | *0.026089* |
| Rps17 | -0.08051 | *>0.999999* | 1.415 | *0.019205* |
| Rps18 | 1.103 | *0.818857* | 1.217 | *0.025668* |
| Rps19 | 0.5216 | *>0.999999* | 1.42 | *0.019205* |
| Rps2 | 0.6832 | *>0.999999* | 1.475 | *0.018026* |
| Rps20 | 0.06227 | *>0.999999* | 1.442 | *0.01879* |
| Rps23 | 0.4285 | *>0.999999* | 1.205 | *0.026243* |
| Rps24 | -0.08688 | *>0.999999* | 1.114 | *0.033162* |
| Rps25 | 0.07025 | *>0.999999* | 1.343 | *0.021333* |
| Rps26 | -0.1308 | *>0.999999* | 1.208 | *0.026089* |
| Rps27a | -0.1596 | *>0.999999* | 0.7906 | *0.08952* |
| Rps27l | 0.1453 | *>0.999999* | 1.333 | *0.021742* |
| Rps3 | 0.2112 | *>0.999999* | 1.379 | *0.020816* |
| Rps3a | 0.2311 | *>0.999999* | 1.305 | *0.022838* |
| Rps4x | 0.6285 | *>0.999999* | 1.254 | *0.02392* |
| Rps5 | 0.2262 | *>0.999999* | 1.753 | *0.008854* |
| Rps6 | 0.2568 | *>0.999999* | 1.252 | *0.023946* |
| Rps7 | 0.09218 | *>0.999999* | 1.36 | *0.021333* |
| Rps8 | 0.1298 | *>0.999999* | 1.249 | *0.024122* |
| Rps9 | 0.5437 | *>0.999999* | 1.406 | *0.019503* |
| Rpsa | 0.1382 | *>0.999999* | 1.22 | *0.025536* |
| Rrbp1 | 0.3912 | *>0.999999* | 1.457 | *0.018271* |
| Ryr1 | -0.6145 | *>0.999999* | 0.4937 | *0.206888* |
| S100a10 | 0.4449 | *>0.999999* | 0.6937 | *0.119262* |
| Sdha | 0.4041 | *>0.999999* | 1.045 | *0.041702* |
| Sdhb | 0.03318 | *>0.999999* | 0.8751 | *0.069052* |
| Sec23a | -0.7851 | *>0.999999* | 1.725 | *0.009403* |
| Sept2 | 0.1586 | *>0.999999* | 1.111 | *0.03321* |
| Sept7 | 0.155 | *>0.999999* | 1.152 | *0.030327* |
| Serpina1c | 1.579 | *0.419802* | 0.2112 | *0.388929* |
| Serpina3n | 1.866 | *0.206398* | 0.9611 | *0.053485* |

| | | | | |
|---|---|---|---|---|
| Serpinf1 | 1.092 | *0.819054* | 1.567 | *0.014388* |
| Serpinh1 | 0.004353 | *>0.999999* | 1.983 | *0.002731* |
| Slc25a12 | -0.3464 | *>0.999999* | 1.17 | *0.029043* |
| Slc25a3 | 0.5334 | *>0.999999* | 1.242 | *0.024265* |
| Slc25a4 | -1.115 | *0.818857* | 0.7915 | *0.08946* |
| Slc25a5 | -0.349 | *>0.999999* | 1.141 | *0.030854* |
| Snd1 | 0.09854 | *>0.999999* | 1.26 | *0.02383* |
| Spon1 | -0.05521 | *>0.999999* | 1.363 | *0.021333* |
| Spta1 | 1.103 | *0.818857* | 1.203 | *0.02636* |
| Sptan1 | 0.1674 | *>0.999999* | 1.029 | *0.043832* |
| Sptb | 0.8509 | *0.962632* | 0.6415 | *0.137912* |
| Sptbn1 | -0.1701 | *>0.999999* | 0.4897 | *0.208728* |
| Srl | -0.7109 | *>0.999999* | 1.349 | *0.021333* |
| Srpx2 | 0.4596 | *>0.999999* | 1.548 | *0.014713* |
| Srsf6 | -0.3155 | *>0.999999* | 1.289 | *0.022838* |
| Ssr4 | 0.2827 | *>0.999999* | 1.206 | *0.026243* |
| Sucla2 | 0.4729 | *>0.999999* | 1.278 | *0.023131* |
| Suclg1 | -0.7036 | *>0.999999* | 1.296 | *0.022838* |
| Synpo2 | 0.06386 | *>0.999999* | 1.155 | *0.030101* |
| Tcp1 | 0.3442 | *>0.999999* | 1.289 | *0.022838* |
| Tf | 2.008 | *0.158741* | 1.247 | *0.024182* |
| Tgfbi | 0.3634 | *>0.999999* | 0.6821 | *0.123155* |
| Thbs1 | 1.257 | *0.709595* | 1.781 | *0.007987* |
| Thbs2 | 0.5264 | *>0.999999* | 2.03 | *0.002642* |
| Thbs3 | 0.2927 | *>0.999999* | 0.4094 | *0.253503* |
| Thbs4 | 1.244 | *0.709595* | 1.952 | *0.00303* |
| Tln1 | 0.09008 | *>0.999999* | 1.399 | *0.019794* |
| Tmod4 | -0.04263 | *>0.999999* | 0.825 | *0.08122* |
| Tnc | 0.09104 | *>0.999999* | 0.9229 | *0.059242* |
| Tnn | -0.5693 | *>0.999999* | 1.111 | *0.03321* |
| Tnnc2 | -0.2314 | *>0.999999* | 1.159 | *0.029986* |
| Tnni2 | -1.449 | *0.505056* | 1.019 | *0.044597* |
| Tnnt3 | -1.889 | *0.196129* | 1.155 | *0.030101* |
| Tpi1 | 0.0847 | *>0.999999* | 0.7865 | *0.090664* |
| Tpm1 | -0.7398 | *>0.999999* | 0.9373 | *0.057515* |
| Tpm2 | -0.4904 | *>0.999999* | 0.5773 | *0.163834* |
| Tpm3 | 0.7812 | *>0.999999* | 0.6735 | *0.126176* |
| Tpt1 | 1.128 | *0.817023* | 0.9526 | *0.054933* |
| Trdn | 1.199 | *0.735812* | 1.06 | *0.039742* |
| Trim72 | -0.806 | *0.995305* | 1.348 | *0.021333* |
| Ttn | -0.4439 | *>0.999999* | 0.7308 | *0.106408* |
| Tuba1c | -0.7989 | *0.998346* | 2.124 | *0.001556* |
| Tuba4a | -0.3844 | *>0.999999* | 0.6254 | *0.143872* |
| Tubb2a | -0.5712 | *>0.999999* | 1.721 | *0.009403* |
| Tubb4b | -0.1845 | *>0.999999* | 1.555 | *0.014388* |
| Tubb5 | -0.09928 | *>0.999999* | 1.992 | *0.002731* |
| Tubb6 | -0.425 | *>0.999999* | 1.997 | *0.002731* |
| Tufm | -0.6464 | *>0.999999* | 1.12 | *0.032739* |
| Uba1 | -0.09428 | *>0.999999* | 1.599 | *0.013906* |
| Ugdh | 0.3376 | *>0.999999* | 1.389 | *0.019857* |
| Ugp2 | 0.4085 | *>0.999999* | 1.239 | *0.024458* |
| Uqcrc1 | 0.735 | *>0.999999* | 0.6801 | *0.123666* |
| Uqcrc2 | 0.1913 | *>0.999999* | 1.11 | *0.03321* |
| Uqcrfs1 | -0.1913 | *>0.999999* | 1.095 | *0.03474* |
| Uqcrq | 0.2015 | *>0.999999* | 0.403 | *0.257147* |
| Vat1 | -0.2079 | *>0.999999* | 1.391 | *0.019857* |
| Vcp | 0.4288 | *>0.999999* | 1.148 | *0.030535* |
| Vdac1 | 0.222 | *>0.999999* | 1.027 | *0.043942* |
| Vdac2 | 0.9014 | *0.950388* | 0.8577 | *0.072452* |
| Vim | -0.4157 | *>0.999999* | 0.8715 | *0.069646* |
| Ywhab | -0.05117 | *>0.999999* | 1.008 | *0.046196* |
| Ywhae | 1.39 | *0.54725* | 1.302 | *0.022838* |
| Ywhag | 0.5833 | *>0.999999* | 0.649 | *0.134877* |
